# Autoantigenic properties of the aminoacyl tRNA synthetase family in idiopathic inflammatory myopathies

**DOI:** 10.1101/2022.06.13.495951

**Authors:** Charlotta Preger, Antonella Notarnicola, Cecilia Hellström, Edvard Wigren, Cátia Fernandes-Cerqueira, Helena Idborg, Ingrid E. Lundberg, Helena Persson, Susanne Gräslund, Per-Johan Jakobsson

## Abstract

**Objectives:** Autoantibodies are thought to play a key role in the pathogenesis of idiopathic inflammatory myopathies (IIM). However, up to 40% of IIM patients, even those with clinical manifestations of anti-synthetase syndrome (ASSD), test seronegative to all known myositis-specific autoantibodies (MSAs). We hypothesized the existence of new potential autoantigens among human cytoplasmic aminoacyl tRNA synthetases (aaRS) in patients with IIM.

**Methods:** Plasma samples and clinical data from 217 patients with, 50 patients with ASSD, 165 without, and two with unknown ASSD status were included retrospectively, as well as serum from 156 age/sex-matched population controls. Samples were screened using a multiplex bead array assay for presence of autoantibodies against a panel of 118 recombinant protein variants, representing 33 myositis-related proteins, including all 19 cytoplasmic aaRS.

**Results:** We identified reactivity towards 16 aaRS in 72 of the 217 patients. Twelve patients displayed reactivity against nine novel aaRS. The novel autoantibody specificities were detected in four patients previously seronegative for MSAs and in eight with previously detected MSAs. We also confirmed reactivity to four of the most common aaRS (Jo1, PL12, PL7, and EJ (n=45)) and identified patients positive for anti-Zo, -KS, and -HA (n=10) that were not previously tested. A low frequency of anti-aaRS autoantibodies was detected in controls.

**Conclusion:** Our results suggest that most, if not all, cytoplasmic aaRS may become autoantigenic. Autoantibodies against new aaRS may be found in plasma of patients previously classified as seronegative with potential high clinical relevance.

## 1. Introduction

Idiopathic inflammatory myopathies (IIM) are characterized by a broad spectrum of clinical manifestations with high mortality and morbidity [1, 2]. Autoantibodies have been identified in more than 50% of patients with IIM, and autoimmunity is thought to play a key role in the pathogenesis of the disease. One sub-group of IIM, named anti-synthetase syndrome (ASSD), is characterized by the presence of autoantibodies targeting aminoacyl transfer(t) RNA synthetases (aaRS), together with specific clinical manifestations such as myositis, interstitial lung disease (ILD), arthritis, mechanic’s hand, Raynaud’s phenomenon, and fever [3, 4].

There are nineteen cytoplasmic aaRS in human cells, including the bifunctional EPRS (Glu-ProRS), one for each amino acid [5]. The most common anti-aaRS autoantibody (anti-Jo1), targeting histidyl tRNA synthetase (HisRS), is present in up to 20-30% of IIM patients [3], and up to 90% of patients with IIM and ILD [6, 7]. Besides HisRS, there are seven other identified autoantigens within the aaRS family in IIM/ASSD [8-11]. Of these, only five are included in the most commonly used commercial assays; anti-Jo1, -PL7, -PL12, -EJ, and -OJ (anti-HisRS, -ThrRS, -AlaRS, -GlyRS, and -IleRS, respectively) [8], indicating a possible underrepresentation of the number of positive patients with anti-aaRS autoantibodies. In addition, there is a potential presence of non-identified anti-aaRS autoantibodies targeting one of the other cytoplasmic aaRS proteins.

A few studies have mentioned additional autoantigens within the human aaRS family, including LysRS (SC), TrpRS (WARS), GlnRS (JS), and SerRS [12-15]. Currently, there is limited data available on the detection of these additional aaRS autoantigens. Moreover, anti-OJ autoantibodies targeting IleRS, one of the members of the intracellular multi-synthetase complex (MSC), have been suggested to potentially target several members of this complex [16, 17], which consists of eight aaRS and three scaffold proteins; aaRS complex interaction proteins (AIMP) 1, -2 and -3 [18].

In this study, we tested the hypothesis that the entire aaRS family displays autoantigenic properties. In addition, we explored the correlations between clinical manifestations and anti-aaRS autoantibody status within patients with ASSD and IIM.

## 2. Materials and methods

### 2.1 Patients and population controls

Plasma samples from 217 consecutive patients with IIM attending Karolinska University Hospital between 1995 and 2014 were retrospectively identified for this cross-sectional study. Classification of IIM was according to the European League Against Rheumatism/American College of Rheumatology (EULAR/ACR) criteria (probability threshold of 55%) [19]. The 2017 European Neuromuscular Centre (ENMC) criteria were applied to classify immune-mediated necrotizing myopathies (IMNM) [20]. Patients were further sub-grouped into ASSD or non-ASSD based on Connors criteria [21], including at least one positive test for any of five anti-aaRS antibodies (anti-Jo1, -PL7, -PL12, -EJ, and -OJ) ever tested by line blot (Euroimmun), immunoprecipitation or ELISA, together with one or more of the following clinical manifestations: ILD, myositis, arthritis, Raynaud’s phenomenon, fever, or mechanic’s hands. Diagnosis of ILD was based on the American Thoracic Society criteria [22]. High-resolution computed tomography (HRCT) and spirometry data were checked for consistent features of ILD. Cardiac involvement was considered if any of the following events occurred during the disease course: pericarditis, myocarditis, arrhythmia, sinus tachycardia. Cancer diagnosis was assigned to patients if ever confirmed during the follow-up (interval between time of diagnosis and last visit at the Rheumatology Clinic). Smoking status was defined as never/ever smoker. Ethnicity was determined at the first visit by the patient self-reporting, and then each patient’s ethnicity has been classified by the responsible physician according to a fixed set of categories. Immunosuppressive treatment was recorded at the time of the plasma sampling. Human leukocyte antigen (HLA)-DRB1 genotyping data was retrieved as previously described [23] for selected patients. For more information see Supplementary Material. The 156 population controls were individuals not affected by rheumatoid arthritis or IIM retrospectively identified from a local biobank, and they were age and sex-matched with the 217 IIM patients on group level (Supplementary Table 1). To control for sample differences between serum and plasma, we compared available plasma and sera from 151 patients with IIM (Supplementary Methods, and Supplementary Fig. 1). This study was approved by the Ethics Committee at Karolinska Institutet, Sweden. All patients gave written informed consent.

### 2.2 Recombinant proteins

Two sets of proteins were used in the multiplex bead array assay. The first set consisted of 25-150 amino acid long protein epitope signature tags (PrESTs), with a median of 100 amino acids and were generated within the Human Protein Atlas (www.proteinatlas.org). The PrESTs are produced in *Escherichia coli* (*E. coli*) and have an affinity tag consisting of a hexahistidine tag and an albumin binding protein domain from streptococcal protein G (His_6_ABP). All PrESTs represents a protein sequence with low homology to other human proteins [24, 25]. The second set of proteins were produced in *E. coli* with an Avi-tag for site-specific biotinylation as previously described [26]. The amino acid coverage was based on clinical interest and solved crystal structures. The selection of antigens used in this study (Supplementary Data) was based on covering the complete human cytoplasmic aaRS protein family, in combination with other known, and available, myositis-specific autoantigens.

### 2.3 Multiplex bead array assay

Neutravidin or PrESTs was amine coupled onto color-coded magnetic beads (Magplex Luminex Corp.) as previously described [27, 28], and internal controls were included. The next day, biotinylated proteins were added to the neutravidin coupled beads and incubated overnight at 4°C. The following day all beads were pooled, and the volume was adjusted to enable the addition of 500 beads per ID to each sample well in a 384 well plate.

Plasma or serum was diluted (1:250) in assay buffer (phosphate buffered saline (PBS), 0.05% (v/v) Tween-20, 3% (w/v) bovine serum albumin (BSA), 0.01 mg/ml neutravidin and 0.16 mg/ml hexahistidine and albumin binding protein tag (His6ABP)) and incubated for 1 h. Beads and diluted plasma or serum were added to each well, and the plate was incubated for 2 h before washing three times with PBS-T (0.05% (v/v) Tween-20). Captured antibodies were fixated to the beads in 0.2 % paraformaldehyde [28] for 10 min before washing three times with PBS-T. Secondary R-Phycoerythrin conjugated Goat F(ab’)2 Fragment anti-Human IgG (γ) (H10104, Invitrogen) was added, the plate was incubated for 30 min before washing three times with PBS-T and final addition of PBS-T to each well. The samples were analyzed on FLEXMAP3D (Luminex Corp.), using xPONENT software (Luminex Corp.), recording median fluorescence intensity (MFI).

### 2.4 ELISA

An ELISA was developed to validate the new anti-aaRS findings. Briefly, biotinylated recombinant proteins were added to streptavidin-coated plates. Plasma was diluted and added before adding a horseradish peroxidase-conjugated anti-human IgG antibody and TMB substrate. For more details see Supplementary Methods.

### 2.5 Statistical analysis

The bead array data were processed in R using RStudio. Based on the quality control analysis, the MFI signals were normalized by antigen in the analysis of serum, scaling the 25^th^ percentile of each antigen to a common value. All samples were normalized per sample by transforming the median fluorescence intensity (MFI) values per sample into number of median absolute deviations (MADs) around the sample median for both sample types [28]. For reproducibility purposes, the multiplex bead array assay was run twice for plasma. Samples that yielded a higher value than the cut-off in both runs, for any of the included versions of the specific protein, were assigned positive. Four different cut-offs were tested, before selecting 100xMAD (Supplementary Table 2, Supplementary Fig. 2-3).

Statistical analyses were performed with Statistical Package for the Social Sciences (SPSS, version 22.0, IBM software, USA). Continuous variables with normal distribution were presented as means with standard deviations (SD), while variables that violated normality were presented as medians with inter-quartile range (IQR). Groups were compared using the independent sample t-test and Mann Whitney U tests. Differences in distributions of categorical variables between groups were tested using the chi-square test and Fisher’s exact test when appropriate. Agreement between the results obtained by different tests was calculated using Cohen’s Kappa coefficient.

Principal component analysis (PCA) for binary data was performed in R using Rstudio (prcomp) to dimensionally reduce the binary data of clinical manifestations. Variables were centered but not scaled. If a patient was positive for the specific manifestation or phenotype, 1 was assigned and 0 was assigned if negative. 140/2449 (5.7%) of the data points were not available (NA). The PCA analysis was done in three different ways assigning NA to either; 0, 1 or randomly 0 or 1, to evaluate the that the NA did not affect the analysis (data not shown), and randomly selected 0 or 1 was used. After analysis, the patients were grouped according to ASSD status.

## 3. Results

### 3.1 IIM cohort: comparison between ASSD and non-ASSD patients

Demographics, laboratory, and clinical data of the IIM cohort (95% Caucasian), comparing 50 patients with ASSD and 165 without ASSD (ASSD status not available for 2/217 patients), is presented in Table 1. Raynaud’s phenomenon, arthritis, ILD, and cardiac disease were statistically more frequent in the ASSD group, while dysphagia was more prevalent among the non-ASSD patients (Table 1). Among myositis-specific autoantibodies (MSAs), anti-Jo1 reactivity was most frequent in the ASSD group, while anti-TIF1*γ* was most common in the non-ASSD group. 69% of patients without ASSD were seronegative for any MSAs.

**Table 1.** Demographic data of the 217 patients with IIM included in the study, 50 with ASSD, 165 without ASSD, and two with unknown ASSD status.

|  | IIM<br>(n=217) | ASSD<br>(n=50) | Non-ASSD<br>(n=165) | p-value* |
| --- | --- | --- | --- | --- |
| Age at diagnosis, mean years (SD) | 56.6 (15.2) | 50.6 (15.6) | 58.3 (14.6) | **0.001** |
| Sex, n (%) women | 137 (63.1) | 34 (68.0) | 101 (61.2) | NS |
| IIM subgroup, n (%) |  |  |  | **0.0001** |
| No myositis | 1 (0.5) | 1 (2.0) | 0 (0.0) |  |
| PM, n (%) | 99 (45.6) | 37 (74.0) | 62 (37.6) |  |
| DM, n (%) | 75 (34.6) | 9 (18.0) | 64 (38.8) |  |
| ADM, n (%) | 5 (2.3) | 1 (2.0) | 4 (2.4) |  |
| sIBM, n (%) | 31 (14.3) | 0 (0.0) | 31 (18.8) |  |
| IMNM**, n (%) | 4 (1.8) | 0 (0.0) | 4 (2.4) |  |
| Ethnicity, n (%) White | 206 (94.9) | 49 (98.0) | 155 (93.9) | NS |
| Disease duration, median years (IQR) | 0 (3) | 0 (3) | 0 (3) | NS |
| Follow-up duration, mean years (SD) | 11.2 (7.8) | 12.4 (8.6) | 11 (7.5) | NS |
| Dead during follow-up, n (%) | 92 (42.4) | 17 (34.0) | 73 (44.2) | NS |
| Age at death, mean (SD) | 75 (10.9) | 73.1 (12.5) | 75.7 (10.6) | NS |
| Autoantibodies |  |  |  |  |
| anti-Jo1, n (%) | 45 (20.7) | 45 (90) | 0 (0.0) | **0.0001** |
| anti-PL7, n (%) | 2 (0.9) | 2 (4.0) | 0 (0.0) | 0.055 |
| anti-PL12, n (%) | 1 (0.5) | 1 (2.0) | 0 (0.0) | NS |
| anti-EJ, n (%) | 1 (0.5) | 1 (2.0) | 0 (0.0) | NS |
| anti-OJ, n (%) | 1 (0.5) | 1 (2.0) | 0 (0.0) | NS |
| anti-Mi-2, n (%) | 5 (2.3) | 0 (0.0) | 5 (3.1) | NS |
| anti-SRP, n (%) | 7 (3.2) | 0 (0.0) | 7 (4.3) | NS |
| anti-MDA5 n (%) | 14 (6.5) | 0 (0.0) | 14 (8.6) | **0.02** |
| anti-TIF1γ, n (%) | 23 (10.6) | 0 (0.0) | 22 (13.6) | **0.003** |
| anti-SSA***, n (%) | 63 (29) | 25 (50.0) | 38 (23.2) | **0.0001** |
| anti-SSB, n (%) | 9 (4.1) | 1 (2.0) | 8 (4.9) | NS |
| anti-U1RNP, n (%) | 19 (8.8) | 6 (12.0) | 13 (7.9) | NS |
| anti-Ku, n (%) | 2 (0.9) | 0 (0.0) | 2 (1.2) | NS |
| anti-Pm-Scl***, n (%) | 20 (9.2) | 3 (6.5) | 17 (10.5) | NS |
| Seronegative (no MSAs), n (%) | 115 (53) | 0 (0.0) | 114 (69.1) | **0.0001** |
| Clinical manifestations |  |  |  |  |
| Other autoimmune disease, n (%) | 45 (20.7) | 10 (20.0) | 35 (21.0) | NS |
| Cancer, n (%) | 59 (27.2) | 9 (18.0) | 48 (29.1) | NS |
| Muscle involvement, n (%) | 216 (99.5) | 49 (98.0) | 165 (100) | NS |
| Myopathic weakness, n (%) | 201 (92.6) | 44 (88.0) | 155 (95.7) | NS |
| Muscle enzyme elevation, n (%) | 198 (91.2) | 45 (91.8) | 151 (94.4) | NS |
| Myopathic EMG, n (%) | 137 (63.1) | 29 (72.5) | 106 (74.6) | NS |
| Pathological muscle biopsy, n (%) | 169 (77.9) | 38 (80.9) | 129 (82.7) | NS |
| Skin involvement, n (%) | 106 (48.8) | 24 (48.0) | 80 (48.5) | NS |
| Raynaud's phenomenon, n (%) | 56 (25.8) | 21 (42.0) | 35 (21.2) | **0.04** |
| Arthritis, n (%) | 56 (25.8) | 29 (58.0) | 27 (17.0) | **0.0001** |
| ILD, n (%) | 69 (31.8) | 39 (79.6) | 30 (18.9) | **0.0001** |
| Cardiac involvement, n (%) | 19 (8.8) | 10 (21.3) | 9 (6.0) | **0.004** |
| Dysphagia, n (%) | 108 (49.8) | 13 (26.5) | 93 (57.4) | **0.0001** |
| Smoking, n (%) | 110 (50.7) | 24 (57.1) | 85 (64.4) | NS |
| Treatment at time of sample, n (%) | 99 (45.6) | 25 (51) | 74 (45.4) | NS |

IIM, idiopathic inflammatory myopathies; ASSD, anti-synthetase syndrome; PM, polymyositis; DM, dermatomyositis; ADM, amyopathic dermatomyositis; sIBM, sporadic inclusion body myositis; IMNM, immune-mediated necrotizing myopathy; Jo1, HisRS; PL7, ThrRS; PL12, AlaRS; EJ, GlyRS; OJ, IleRS; Mi-2, chromatin organization modifier helicase (CHD) 3 and 4; SRP, signal recognition particle; MDA5, interferon-induced helicase C domain-containing protein 1; TIF1 $\gamma$ , E3 ubiquitin-protein ligase TRIM33; SSA, Ro52 (tripartite motif containing 21 (TRIM21)) and Ro60 (TROVE domain family member 2); SSB, Sjogren syndrome antigen B; U1RNP, small nuclear ribonucleoprotein U1 subunit 70; Ku, X-ray repair cross complementing (XRCC) 6; Pm-Scl, polymyositis-scleroderma overlap syndrome-associated antigen 75 (exosome component 9) and 100 (exosome component 10); MSA, myositis-specific autoantibodies; EMG, electromyography; ILD, interstitial lung disease.
Disease duration = interval between the time of diagnosis and the time of sampling; follow-up duration = interval between the time of diagnosis and the time of the last recorded visit at the Rheumatology Unit,
Karolinska University Hospital, Sweden. Patients with unknown data were not included in the table nor the comparison for each information. \* the reported p-value is for comparisons between the ASSD and the non-ASSD group of patients (information on ASSD status was not available for two patients, excluded from the two groups). \*\* all patients with IMNM tested positive for anti-HMGCR antibodies. \*\*\* with regards to anti-SSA antibodies, information on reactivity to the individual Ro52 (TRIM21) or Ro60 (TROVE2) was not available for all patients and therefore not reported. The same applied to Pm-Scl where the commercial test included both Pm-Scl 75 and Pm-Scl 100 but information regarding the separate antigens was not available.

### 3.2 Autoantibodies detected in the multiplex bead array assay

In the IIM cohort, autoantibodies against all cytoplasmic aaRS proteins except three (IleRS (OJ), LeuRS, and AspRS) were detected (Fig. 1). Autoantibodies against any of the aaRS were present in one-third (n=72, 33%), and of these, seven patients were positive for two and one patient for three anti-aaRS antibodies (Supplementary Table 3). Nine patients from the non-ASSD group were positive for anti-Jo1, -PL7, -PL12, or -EJ (Supplementary Table 4). In addition, we detected reactivities to other MSA antigens (MDA5, Mi-2, and TIF1*γ*), myositis-associated autoantibody (MAA) antigens (SSA (Ro52 (TRIM21)), SSB, U1RNP, and Pm-Scl), and to AIMP-1 and AIMP-2, two of the MSC scaffold proteins (Fig. 1 and Fig. 2D, Supplementary Table 5).

**Fig 1.**
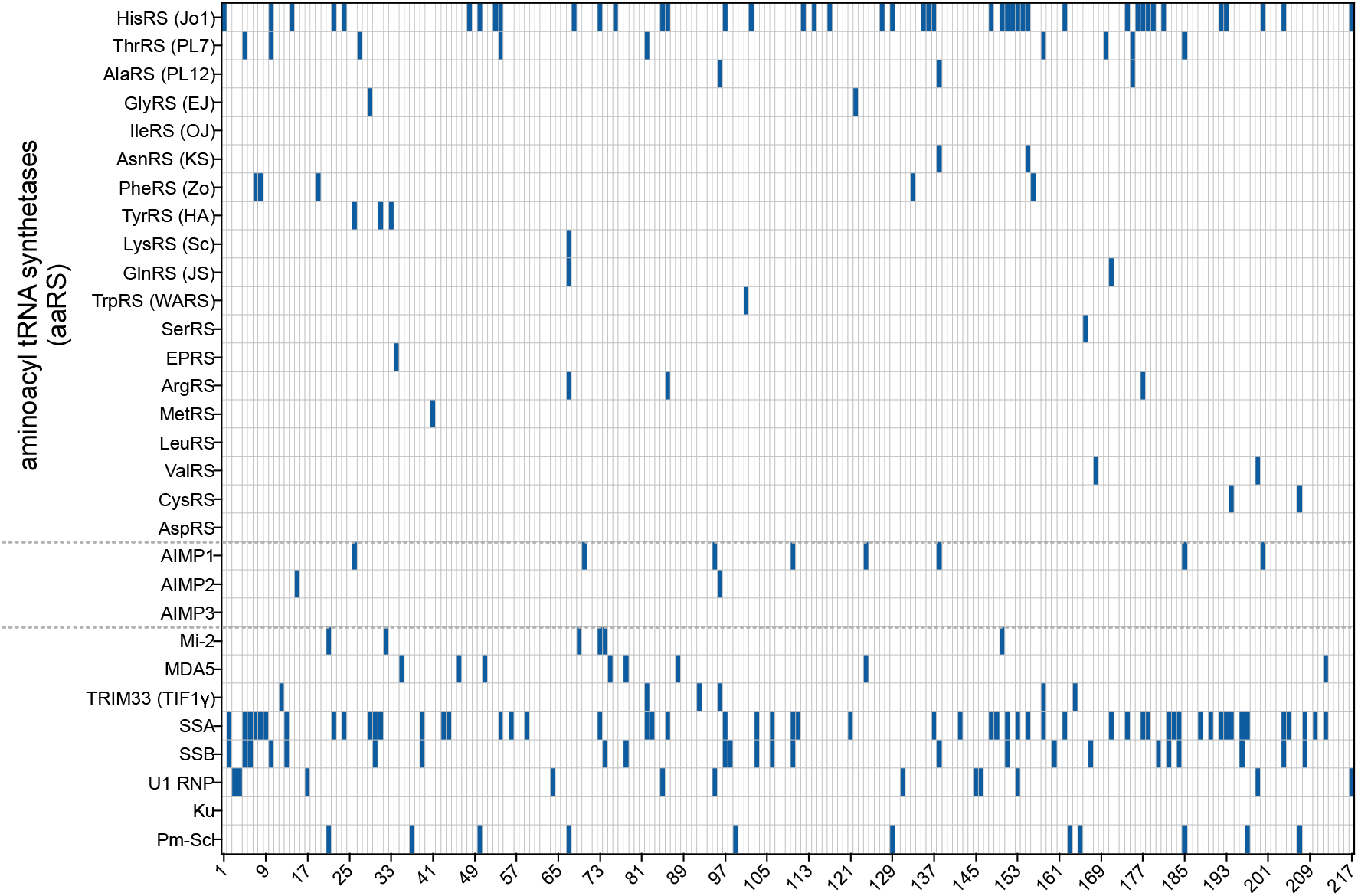
Autoantibody reactivities for IIM patients. Reactivity against a panel of 30 antigens for 217 IIM patients as assessed by the multiplex bead array assay. Each column represents one patient, (patient 1-217), and each row represents one potential autoantigen. Reactivity was assigned positive (blue) if the criteria as defined in the method section were met for at least one of the included versions of a particular protein antigen. All cytoplasmic aaRS proteins are displayed above the dotted gray lines, the AIMP proteins are in between dotted lines, and below are the additional myositis-related proteins included in the study. With regards to anti-SSA reactivities, all 56 IIM positive patients were reactive against Ro52 (TRIM21) and none against Ro60 (TROVE2), using 100xMADs as a cut-off.

**Fig 2.**
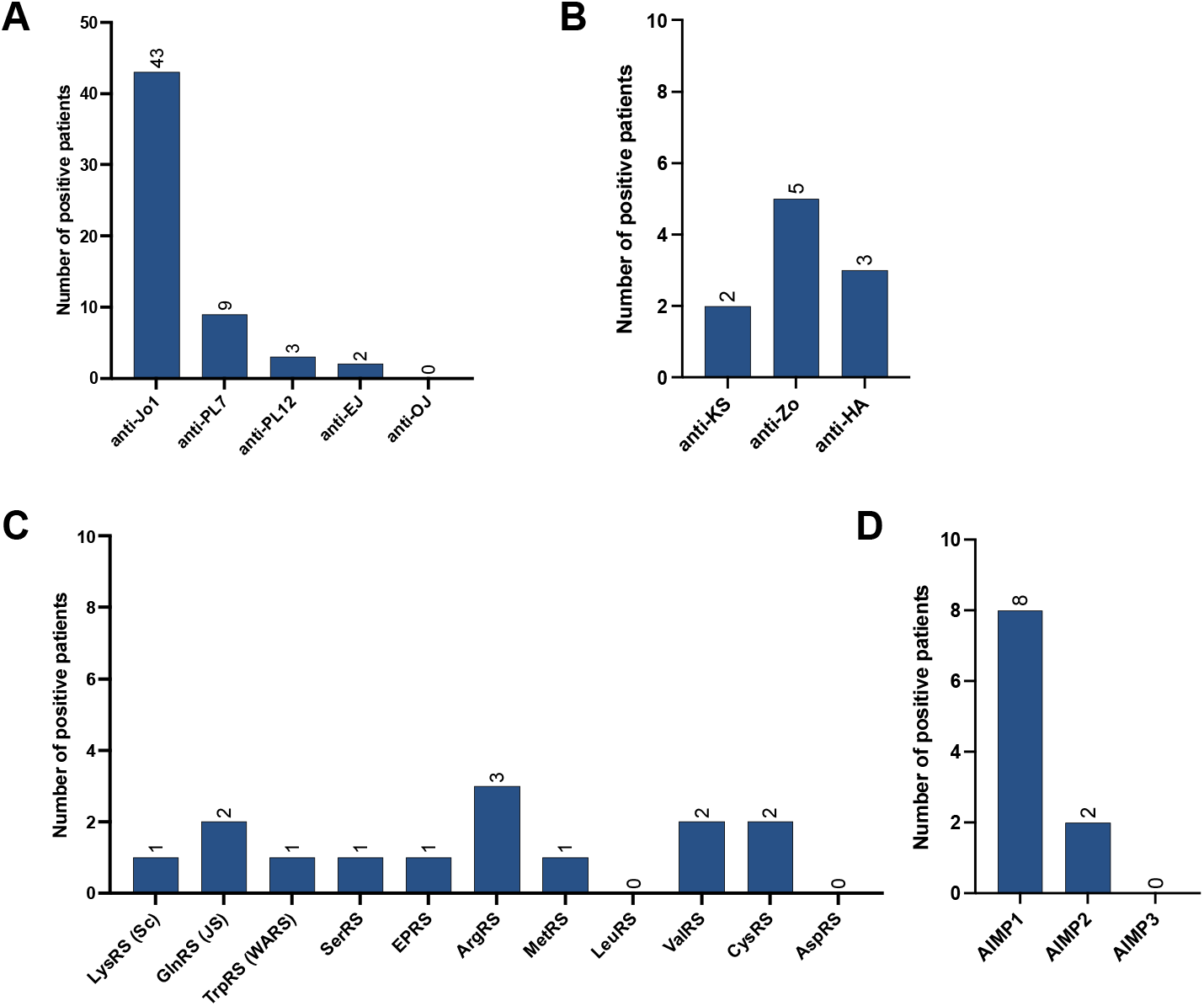
IIM patients positive for autoantibodies against aaRS and AIMPs using the multiplex bead array assay. Patients with autoantibodies targeting; **(A)** the five aaRS autoantigens usually tested for in the clinic, **(B)** known ASSD-associated aaRS autoantigens usually not tested for in clinical settings, and **(C)** the remaining eleven human cytoplasmic aaRS not previously associated to IIM/ASSD as autoantigens. **(D)** Patients positive for AIMP (1-3), the three scaffold proteins that are part of the multi-synthetase complex (MSC).

Autoantibodies towards nine aaRS (LysRS, GlnRS, TrpRS, SerRS, EPRS, ArgRS, MetRS, ValRS, and CysRS), not previously associated with IIM/ASSD were detected in 12 patients (Fig. 2C, Table 2). Of these, four were in the seronegative group, *i.e*., not presenting any other MSAs, while eight had previously tested positive for MSAs (anti-Jo1 (n=3), -MDA5 (n=2), - Mi2 in combination with -TIF1*γ* (n=1) -TIF1*γ* (n=1) and -SRP (n=1), Table 2). Of these eight, we could confirm anti-Jo1 autoantibodies in two of three patients but not the other previously reported MSAs (Table 2). Reactivities to known aaRS autoantigens in ASSD, not previously tested in this cohort, were found in 10 individuals: AsnRS (KS, n=2), PheRS (Zo, n=5), and TyrRS (HA, n=3) (Fig. 2B, Table 2). In addition, we identified patients with multiple reactivities, both with known and potential novel anti-aaRS as well as other MSAs (Supplementary Table 3).

**Table 2.** Brief characteristics of the patients with IIM who were positive for the new aaRS autoantibody specificities not previously tested in this cohort. **Upper part:** IIM patients (n=12) testing positive for anti-aaRS autoantibodies other than the eight usually described. **Lower part:** IIM patients (n=10) testing positive for autoantibodies anti-KS, -HA, and -Zo in this study. Previously known autoantibody status, smoking status, and clinical manifestations are included. The autoantigen for the specific autoantibody is stated in the table.

| Patients with IIM (n=12) positive for novel anti-aaRS autoantibodies |  |  |  |  |  |  |  |
| --- | --- | --- | --- | --- | --- | --- | --- |
| Patient | Clinical subgroup | Known antibody positivity | aaRS detected in this study | Smoking status | HLA-DRB1 Allele1/Allele2 | Clinical manifestations | Validated by ELISA |
| 34 | non-ASSD | seroneg | EPRS | yes | *03/*15 | PM | Suppl. Fig. 6 |
| 41 | non-ASSD | seroneg | MetRS | yes | *03/*07 | PM | Fig. 4 |
| 67 | non-ASSD | seroneg | LysRS, GlnRS and ArgRS | yes | *03/*04 | DM, mechanic's hands | Suppl. Fig. 7 |
| 86 | ASSD | HisRS (Jo1) | HisRS (Jo1) and ArgRS | na | *03/*09 | DM, mechanic's hands, ILD, arthritis, cancer |  |
| 101 | non-ASSD | MDA5 | TrpRS (WARS) | na | *03/*04 | PM, arthritis | Suppl. Fig. 6 |
| 166 | non-ASSD | Mi-2 and TIF1g | SerRS | na | *04/*15 | DM, ILD, DM skin features, dysphagia, cancer | Suppl. Fig. 6 |
| 168 | ASSD | HisRS (Jo1) | ValRS | yes | *08/*14 | PM | Fig. 4 |
| 171 | non-ASSD | MDA5 | GlnRS | yes | *01/*03 | PM |  |
| 177 | ASSD | HisRS (Jo1) | HisRS (Jo1) and ArgRS | yes | na | Muscle weakness<sup>+</sup>, ILD |  |
| 194 | non-ASSD | TIF1g | CysRS | yes | *01/*03 | DM, cancer | Suppl. Fig. 6 |
| 199 | non-ASSD | SRP<sup>++</sup> | ValRS | yes | *04/*15 | PM |  |

| 207 | non-ASSD | seroneg | CysRS | no | *03/*13 | PM |
| --- | --- | --- | --- | --- | --- | --- |
| **Patients with IIM (n=10) positive for anti-aaRS autoantibodies previously not tested for** |  |  |  |  |  |  |
| **Patient** | **Clinical subgroup** | **Known antibody positivity** | **aaRS detected in this study** | **Smoking status** | **Clinical manifestations** | **Validated by ELISA** |
| 7 | non-ASSD | seroneg | PheRS (Zo) | yes | PM |  |
| 8 | non-ASSD | seroneg | PheRS (Zo) | yes | PM, ILD | Suppl. Fig. 8 |
| 19 | non-ASSD | seroneg | PheRS (Zo) | yes | PM, Raynaud's, dysphagia |  |
| 26 | non-ASSD | seroneg | TyrRS (HA) | na | sIBM, Raynaud's |  |
| 31 | non-ASSD | seroneg | TyrRS (HA) | yes | DM, DM skin features, arthritis, Raynaud's, dysphagia |  |
| 33 | non-ASSD | TIF1g and HGMCR | TyrRS (HA) | no | DM, DM skin features, ILD, Raynaud's, cancer, dysphagia | Suppl. Fig. 8 |
| 133 | non-ASSD | TIF1g | PheRS (Zo) | yes | DM, DM skin features, cardiac involvement, dysphagia, cancer |  |
| 138 | non-ASSD | seroneg | AlaRS (PL12) and AsnRS (KS) | yes | PM, calcinosis, dysphagia | Suppl. Fig. 10 |
| 155 | ASSD | HisRS (Jo1) | HisRS (Jo1) and AsnRS (KS) | yes | DM, DM skin features, ILD, cancer, dysphagia |  |
| 156 | non-ASSD | SRP<sup>++</sup> | PheRS (Zo) | yes | DM, DM skin features, calcinosis, |  |

**Upper part:** Seven of these twelve patients were selected for validation using ELISA and the result are shown in the figure stated in the last column. **Lower part:** Three of the ten patients were selected for validation using ELISA and the results are shown in the figure stated in the last column. HLA-DRB1 data was only included for the 12 IIM patients in the upper part of the table. aaRS, aminoacyl tRNA synthetase; ASSD, anti-synthetase syndrome; PM, polymyositis; DM, dermatomyositis; HMGCR, 3-hydroxy-3-methylglutaryl-coenzyme A reductase; ILD, interstitial lung disease; MDA5, interferon-induced helicase C domain-containing protein 1; Mi-2, chromatin organization modifier helicase (CHD) 3 and 4; sIBM, sporadic inclusion body myositis; na, not available; seroneg, previously no myositis specific autoantibodies detected; Suppl., supplementary; SRP, signal
recognition particle; TIF1 $\gamma$ , E3 ubiquitin-protein ligase TRIM33. Smoking status = yes (ever smoker,) no (never smoker). DM skin features = any of periungual erythema, Gottron's sign, Gottron's papules, V-sign, shawl sign, erythroderma, periorbital edema, heliotrope rash. <sup>+</sup>muscle weakness based on manual muscle test-8 (MMT-8) below 80 and/or impaired muscle endurance by myositis functional index-2; patient 177 did not fulfill European League Against Rheumatism/American College of Rheumatology (EULAR/ACR) criteria for the classification of idiopathic inflammatory myopathies. <sup>++</sup>SRP was not included in the multiplex bead array assay.

In the population controls (PC), 32/156 (20.5%) displayed reactivity against any of the included antigens (Supplementary Fig. 4), and 15 (9.6%) individuals were reactive to any of the nineteen aaRS, with the highest frequency of HA (n=4), ArgRS (n=3), CysRS (n=2), and LeuRS (n=2) (Table 3, Supplementary Fig. 4). Of the nine novel anti-aaRS reactivities found in the IIM cohort, we detected reactivity in PC against five: LysRS (n=1), SerRS (n=1), EPRS (n=1), ArgRS (n=3), CysRS (n=2). Reactivity against Mi-2 (n=9), MDA5 (n=2), SSA (n=5), SSB (n=5) and Pm-Scl (n=2) was also detected. To control for sample discrepancies, 151/217 patients with IIM were analyzed using both serum and plasma, and 134/151 (89%) agreed (Supplementary Fig. 5).

**Table 3.** Number of individuals with reactivity against aaRS in 217 IIM and 156 PC. The autoantigen for the specific autoantibody is stated in the table.

| Antigen | Reactive 217 IIM<br>n (%) | Reactive 156 PC<br>n (%) |
| --- | --- | --- |
| HisRS (Jo1) | 43 (19.8) | 1 (0.6) |
| ThrRS (PL7) | 9 (4.1) | 0 (0.0) |
| AlaRS (PL12) | 3 (1.4) | 0 (0.0) |
| GlyRS (EJ) | 2 (0.9) | 0 (0.0) |
| IleRS (OJ) | 0 (0.0) | 0 (0.0) |
| AsnRS (KS) | 2 (0.9) | 1 (0.6) |
| PheRS (Zo) | 5 (2.3) | 0 (0.0) |
| TyrRS (HA) | 3 (1.4) | 4 (2.6) |
| LysRS (Sc) | 1 (0.5) | 1 (0.6) |
| GlnRS (JS) | 2 (0.9) | 0 (0.0) |
| TrpRS (WARS) | 1 (0.5) | 0 (0.0) |
| SerRS | 1 (0.5) | 1 (0.6) |
| EPRS | 1 (0.5) | 1 (0.6) |
| ArgRS | 3 (1.4) | 3 (1.9) |
| MetRS | 1 (0.5) | 0 (0.0) |
| LeuRS | 0 (0.0) | 2 (1.3) |
| ValRS | 2 (0.9) | 0 (0.0) |
| CysRS | 2 (0.9) | 2 (1.3) |
| AspRS | 0 (0.0) | 0 (0.0) |

aaRS, aminoacyl tRNA synthetase; IIM, Idiopathic inflammatory myopathies; PC, population control.

Clinical manifestations of the 22 patients with autoantibodies against novel aaRS and previously not tested aaRS are summarized in Table 2. Myositis was diagnosed in all patients with anti-HA, -Zo, or -KS (n=10), while ILD affected three. Arthritis was reported by one patient with anti-HA antibodies. The three anti-HA and one anti-Zo positive patients had Raynaud’s phenomenon. None presented with mechanic’s hands. All patients with novel aaRS (n=12) had either muscle weakness and/or muscle enzyme elevation. Electromyography showed myopathic changes in 7/9 patients and muscle biopsy was consistent with myositis in 8/12 patients. Out of eight patients with pathological muscle biopsy, all presented widespread up-regulation of major histocompatibility complex class I (MHC-I), five with perivascular and/or endomysial inflammatory infiltrates even invading non-necrotic muscle fibers and two with perifascicular atrophy. The patient with anti-SerRS antibody reactivity had perifascicular necrosis, which has been proposed to be specific for ASSD [29-31] (missing information in 5/8 pathological muscle biopsies). None of the patients suffered from Raynaud’s phenomenon. Two of the three patients with ILD, of which one also displayed arthritis and mechanic’s hands, had previously tested positive for anti-Jo1 autoantibodies. Of the 12 patients with novel anti-aaRS, only three had negative anti-nuclear antibodies (ANA) by indirect immunofluorescence (IIF), while six presented with ANA positivity and homogeneous, nucleolar, or granular pattern (information not available for three patients).

According to Connors criteria [21], the 22 IIM patients described above could be re-classified as having ASSD. After including these in the previous classified ASSD group, we ended up with 68 patients with ASSD and 147 with non-ASSD. The frequencies of clinical manifestations in the two groups were the same as in the analysis reported in Table 1. Principal component analysis of the clinical manifestations did not show any clear differentiation between the two groups, and the 22 patients with newly detected anti-aaRS reactivities were closer to the non-ASSD group (Fig. 3).

**Fig. 3.**
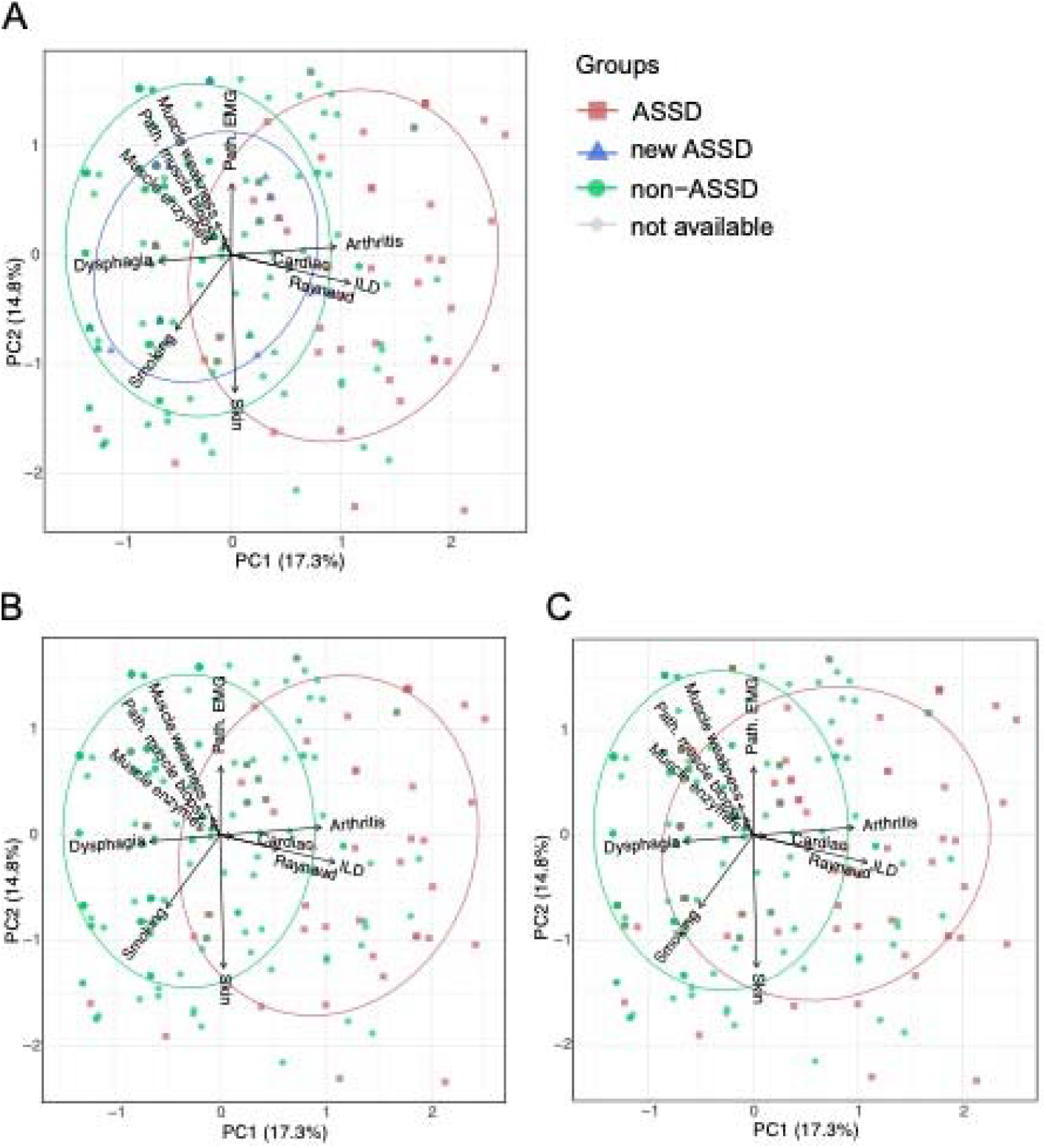
Principal component analysis (PCA) of clinical manifestations and phenotypes. Analysis based on the binary data of the variables; muscle involvements (pathological muscle biopsy, muscle enzymes elevation, pathological EMG, and muscle weakness), skin involvement, Raynaud’s phenomenon, Arthritis, interstitial lung disease (ILD), cardiac involvement, dysphagia, and smoking. Scores plots PC1 vs PC2 are shown, each dot represents one patient and the contribution of each variable to PC1 and PC2 are included. Some dots are overlapping represented by the change of color intensity. Grouping is based on **(A)** ASSD classification, ASSD (n=50, red), non-ASSD (n=147, green), not available ASSD status (n=2, gray), patients with a new ASSD classification after this study (n=18, blue). **(B)** Patients are grouped based on ASSD status from clinical information (before this study) ASSD (n=50, red), non-ASSD (n=165, green), and not available ASSD status (n=2, gray). **(C)** Patents grouped based on ASSD status after reclassifying 22 patients into the ASSD group ASSD (n=68, red), non-ASSD (n=147, green), and not available ASSD status (n=2, gray). Demographic data is according to Table 1. The analysis indicates no clear differentiation between groups in scores plot of PC1 vs PC2.

### 3.3 Measurement of agreement

The antibody results obtained from this study were compared with those previously known and used to stratify the IIM cohort in ASSD and non-ASSD groups (Supplementary Fig. 2-3) by calculating the kappa coefficient (Table 4). 45/50 previously known anti-synthetase autoantibodies could be detected in this study, all except for four anti-Jo1 and one anti-OJ (Supplementary Fig. 2).

**Table 4.** Measurement of agreement between this study and previously known antibody status. The comparison was done using the total number of positive patients for myositis-specific autoantibodies and Cohen’s kappa coefficient.

|  | Known positivity, n | Positivity detected in current study, n | Kappa coefficient | p-value |
| --- | --- | --- | --- | --- |
| Anti-Jo1 | 45 | 43 | 0.91 | 0.0001 |
| Anti-PL7 | 2 | 9 | 0.39 | 0.0001 |
| Anti-PL12 | 1 | 3 | 0.49 | 0.014 |
| Anti-EJ | 1 | 2 | 0.66 | 0.0001 |
| Anti-OJ | 1 | 0 | / | / |
| Anti-Mi-2 | 5 | 6 | -0.027 | NS |
| Anti-MDA5 | 14 | 8 | 0.52 | 0.0001 |
| Anti-TIF1γ | 23 | 6 | 0.31 | 0.0001 |

Jo1, HisRS; PL7, ThrRS; PL12, AlaRS; EJ, GlyRS; OJ, IleRS; Mi-2, chromatin organization modifier helicase (CHD) 3 and 4; SRP, signal recognition particle; MDA5, interferon-induced helicase C domain-containing protein 1; TIF1 $\gamma$ , E3 ubiquitin-protein ligase TRIM33.

### 3.4 ELISA validation

To validate the findings of new anti-aaRS autoantibody reactivities in IIM, one patient representing each new autoantigen was selected, and an ELISA method was developed. We could confirm all but one (GlnRS) autoantibody reactivity (Fig. 4 and Supplementary Fig. 6-11).

**Fig. 4.**
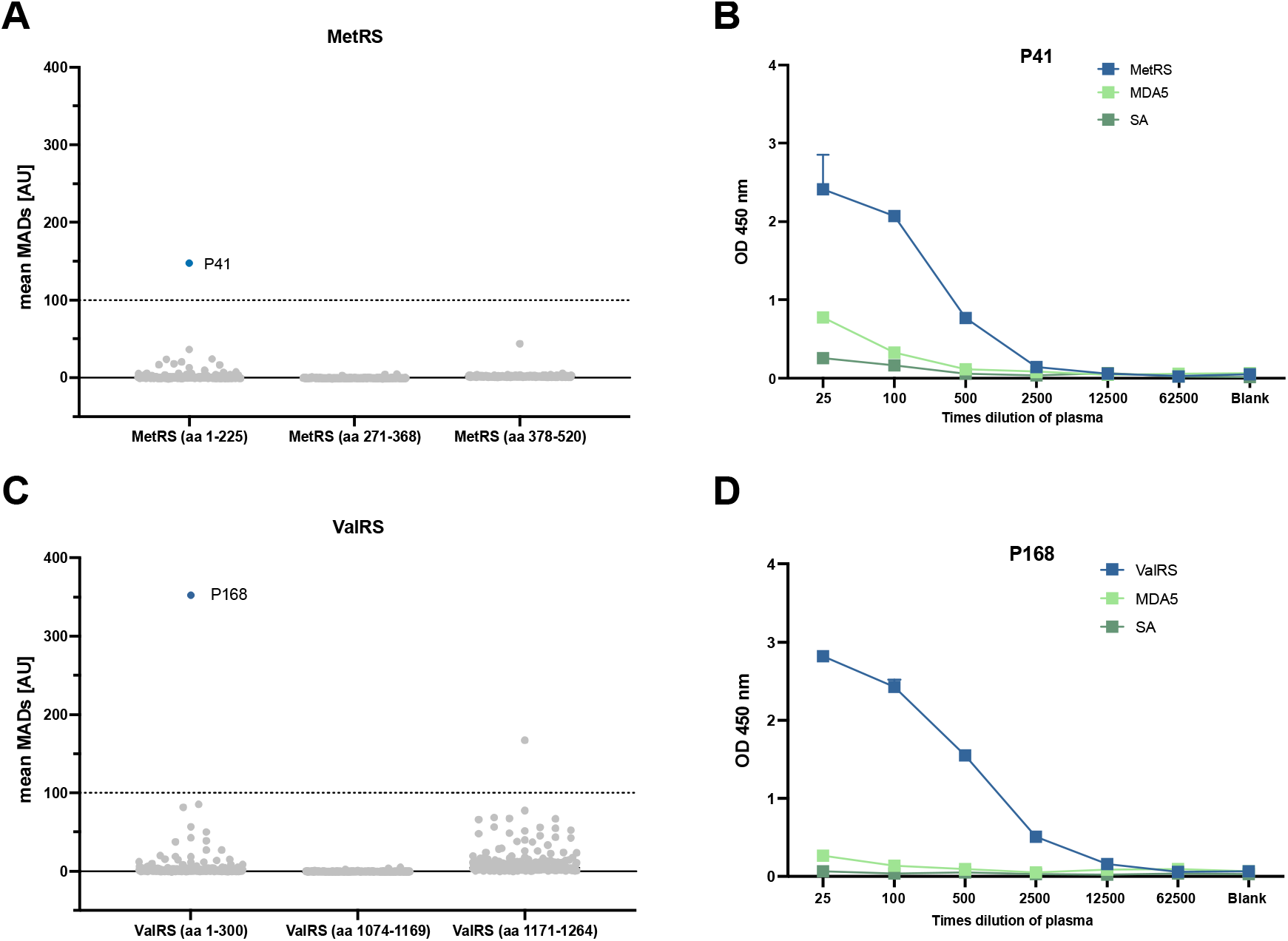
Validation of bead array assay results with ELISA. Mean MADs from the bead array assay showing patients with reactivity against (**A**) MetRS and (**C**) ValRS. Patient (P) 41 showed reactivity against the N-terminal part of MetRS (aa 1-225) and P168 against the N-terminal part of ValRS (aa 1-300) (blue). The distribution of the other 216 patients for each antigen is shown in gray. The dotted gray line represents the cut-off at 100xMADs. Antibody reactivity against MetRS (aa 1-225) and ValRS (aa 1-300) were measured by ELISA and absorbance values (450 nm) obtained for (**B**) P41 and (**D**) P168 are shown (blue). MDA5 (light green) was used as a control protein and streptavidin (SA, green) represents the background signal. A plasma sample from an MDA5 positive patient was used as a control for protein-specific background (Supplementary Fig. 11). The plasma samples were diluted in a five-fold dilution series from 25 times to 62500 times. MADs, median absolute deviations; OD, optical density; P, patient; SA, streptavidin.

## 4 Discussion

In this study, a well-characterized IIM cohort and population controls were screened for autoantibody reactivities against the entire family of cytoplasmic aminoacyl-tRNA synthetases (aaRS). Our results indicate that all cytoplasmic aaRS but two display autoantigenic properties in patients with IIM.

Myositis-specific autoantibodies (MSAs) represent a fundamental diagnostic tool, helping to identify different IIM subgroups characterized by distinct clinical manifestations and histopathological features as well as to predict disease prognosis [32]. However, more than 40% of IIM patients test negative for the commonly tested, generally described MSAs [10], indicating a possibility to identify yet unknown autoantigens.

Here, we explored if patients with IIM test positive for autoantibodies against any of the cytoplasmic aaRS, using a multiplex bead array assay. To increase the possibility of detecting new autoantigens, we included different versions of the same aaRS, either full-length or truncated versions, to allow for detection of autoantibodies targeting both conformational dependent and -independent epitopes. We found that more than one-third of the IIM cohort tested positive for any anti-aaRS antibody, independently of previous autoantibody status. We could detect autoantibodies against 16/19 cytoplasmic aaRS, including nine aaRS proteins that, to our knowledge, have never been described as autoantigens in IIM before or have only been reported in occasional individuals [12-15]. Importantly, reactivities against these novel proteins were identified in patients previously classified as seronegative for MSAs.

For anti-Jo1, -PL12, -PL7, -EJ, and -OJ, we could confirm previously known anti-aaRS antibodies in 45/50 patients, missing only four anti-Jo1 and one anti-OJ reactivities. The low kappa coefficient for anti-PL12, -PL7, and -EJ could be explained by new reactivities found in this study, not previously detected, or tested for. As explained above, the inclusion of several antigens from the same protein might increase the possibility to detect autoantibodies. Moreover, limitations with conventional methods used in the clinic have been noted. For example, anti-aaRS antibodies may be negative in line blot [33], but can show a cytoplasmic ANA pattern by IIF as aaRS are located mainly in the cytoplasm [5, 34].

Thirteen patients had co-existence of anti-aaRS antibodies, or anti-aaRS antibodies together with other MSAs. This is of particular interest as anti-aaRS autoantibodies are usually described as mutually exclusive [8-11]. Since the sequence similarities between the aaRS proteins are low (Supplementary Table 3), it is unlikely that the multiple reactions are due to cross-reactivity [35, 36]. Nevertheless, studies have suggested that autoantibodies from the same individual could target several members of the multi-synthetase complex (MSC) [16, 17, 37]. Here, we found one patient, P67, with autoantibodies targeting three MSC members (ArgRS, GlnRS, and LysRS) corroborating this hypothesis.

There are, to our knowledge, only a limited number of studies available investigating the presence of anti-aaRS autoantibodies in population controls, particularly regarding the rarer anti-aaRS autoantibodies [38-40]. Our study gives additional insight into this. Autoantibodies targeting aaRS and other autoantigens were observed at low frequencies, as expected in control cohorts [41]. However, the relatively high frequency of reactive subjects in the PC with the rarer anti-aaRS was a surprise (Table 3). The fact that we used population controls that might have other autoimmune diseases could explain some of the reactivities. Recent studies reported a relatively high prevalence of anti-Zo, -KS, and -HA in a broad spectrum of ILD patterns [42], and ILD has been reported as the primary clinical feature of anti-KS patients [43]. In our cohort, ten patients with IIM were identified with these autoantibodies, and ILD was reported only in three. Patient selection, in our study from a rheumatology clinic, may explain these differences. Notably, anti-HA antibodies were found at a higher frequency in PC than in the IIM cohort (2.6 vs. 1.4%). The exact meaning of this result needs further investigation, and the low frequencies of the rare anti-aaRS autoantibodies found in both IIM and PC should be further validated in larger cohorts. Still, our study highlights the importance of including population controls in research, but also in clinical routines to define appropriate cut-offs.

Twelve patients were identified with new anti-aaRS autoantibodies. Two-thirds of these were HLA-DRB1*03 positive and current or previous smokers, in line with the known association between HLA-DRB1*03 haplotype, smoking, and anti-aaRS antibodies [44-47]. The ANA-positivity, without cytoplasmic pattern, reported in 6/12 patients could be explained by the co-existents of other MSA or MAA. When investigating the clinical and histopathological features of the 12 patients with novel anti-aaRS autoantibodies, we could not verify the typical characteristics of ASSD, neither in clinical nor histopathological features. However, this small group of patients and the fact that five of these autoantibodies were also found in PC, makes it difficult to draw conclusions regarding their potential association with ASSD. Similarly, anti-TrpRS autoantibodies, although previously detected in patients with autoimmune diseases,[14] have not been suggested as a serological marker for ASSD since the related clinical phenotype was more similar to rheumatoid arthritis than ASSD [16, 48]. Nevertheless, all IIM patients with novel anti-aaRS antibodies presented with muscle involvement.

The novel anti-aaRS autoantibodies were mostly found in the non-ASSD group and in four who were previously known as seronegative. Even though some co-existence of anti-aaRS autoantibodies was found, the majority of anti-aaRS positive individuals only had one detectable anti-aaRS autoantibody. For individuals previously known as seropositive, with novel anti-aaRS autoantibodies detected here (n=8), the previous autoantibody positivity could only be verified in two individuals. The possible reason for these discrepancies are discussed in the paragraph below.

The limitations of this study include the following. Firstly, with the study design used here, it is not possible to conclude if the novel aaRS autoantigens are specific for IIM or not. Both TrpRS and SerRS have previously been suggested as autoantigens in other diseases [13, 14]. Also, the new reactivities were detected in a low frequency in IIM patients, and some also in controls, and confirmation in larger cohorts is needed. Secondly, some samples were retrieved after the patient started immune-modulating treatment, which could affect the presence and detection of autoantibodies [49, 50]. Thirdly, we did not cover the full-length protein of all autoantigens, indicating that we may have some false negatives. For example, anti-OJ reactivity in patient P95 could not be confirmed in this study, in which only shorter protein versions of IleRS were included. Fourthly, sample collection did not always match the timepoint for MSA detection in clinic, and for some patients, data were missing. This could explain why some patients presented discordant results. Finally, to minimize the risk of false positives, we decided to use a high cut-off for all antigens, even though this means a higher risk for false negatives.

In conclusion, our results suggest autoantigenic properties for the cytoplasmic aaRS family, as well as the AIMP proteins, and we hypothesize that in a larger cohort, all aaRS might be found autoantigenic. However, to infer how specific these novel autoantibodies are for IIM, or for distinct clinical phenotypes, these results need to be tested in another large study. There are still remaining seronegative patients left in our cohort, and we suggest to use more multiplex assays in research comprising additional proteins to explore and investigate new potential autoantigens. Combining serological, clinical, and histopathological findings makes it possible to define more homogeneous groups in IIM to achieve an improved understanding of the pathophysiology behind the muscular and extra-muscular manifestations and aim at a more personalized treatment. Here, we also found low frequencies of the novel and previously described anti-aaRS autoantibodies in population controls. For several of the anti-aaRS autoantibodies, frequencies were similar between IIM patients and controls, and this study emphasizes the importance to include population controls in screening for new autoantibodies.

## Acknowledgements

We thank Julia Norkko for the help with sample collection from the biobank. We also thank the SciLifeLab facilities Autoimmunity and Serology Profiling and Human Antibody Therapeutics (Drug Discovery and Development) for experimental assistance and use of their instrumentation and infrastructure.

## Abbreviations

aaRS: aminoacyl transfer (t) RNA synthetase(s)
ADM: amyopathic dermatomyositis
ANA: anti-nuclear antibodies
ASSD: anti-synthetase syndrome
BSA: bovine serum albumin
DM: dermatomyositis
EMG: electromyography
ENMC: European Neuromuscular Centre
EULAR/ACR: European League Against Rheumatism/American College of Rheumatology
His_6_ABP: hexahistidine and albumin binding protein tag
HLA: human leukocyte antigen
HRTC: high-resolution computed tomography
IIF: indirect immunofluorescence
IIM: idiopathic inflammatory myopathies
ILD: interstitial lung disease
IQR: inter-quartile range
IMNM: immune-mediated necrotizing myopathy
MAA: myositis-associated autoantibodies
MADs: median absolute deviations
MFI: median fluorescence intensity
MHC: major histocompatibility complex
MSA: myositis-specific autoantibodies
MSC: multi-synthetase complex
PBS: phosphate buffered saline
PC: population control
PCA: principal component analysis
PM: polymyositis
PrEST: protein epitope signature tag
sIBM: sporadic inclusion body myositis
SD: standard deviation.

A list of all proteins used in this study and abbreviations thereof are available in Supplementary Data.

## Author contributions

All authors have read and approved on the final version to be published

Charlotta Preger: Conceptualization, Data Curation, Methodology, Investigation, Formal analysis, Visualization, Writing – Original draft, Writing – Review & Editing. Antonella Notarnicola: Conceptualization, Data Curation, Investigation, Formal analysis, Visualization, Writing – Original draft, Writing – Review & Editing. Cecilia Hellström: Data Curation, Methodology, Investigation, Formal analysis, Writing – Review & Editing. Edvard Wigren: Investigation, Formal analysis, Writing - Review & Editing. Cátia Fernandes-Cerqueira: Conceptualization, Writing – Review & Editing. Helena Idborg: Formal analysis, Writing – Review & Editing. Ingrid E. Lundberg: Conceptualization, Investigation, Funding acquisition, Supervision, Resources, Writing – Review & Editing. Helena Persson: Conceptualization, Resources, Supervision, Writing – Review & Editing. Susanne Gräslund: Conceptualization, Funding acquisition, Resources, Supervision, Writing – Review & Editing. Per-Johan Jakobsson: Conceptualization, Supervision, Funding acquisition, Writing – Review & Editing.

## Data Availability

The data undelaying this article will be made available on request.

## Disclaimer

This communication reflects the views of the authors and neither IMI nor the European Union, EFPIA or any Associated Partners are liable for any use that may be made of the information contained herein.

