## Supplementary Data for "Autoantigenic properties of the aminoacyl tRNA synthetase family in idiopathic inflammatory myopathies"

| Protein | Protein short name | Gene | Clinical Name | Corresponding autoantibody group | amino acid coverage | amino acid sequence | Uniprot ID | Antigen name | Type |
|---|---|---|---|---|---|---|---|---|---|
| alanyl-tRNA synthetase | AlaRS | AARS1 | PL12 | MSA | 1-455 | MDSTLTASEIRQRFIDFFKRNEHTYVHSSATIPLDDPTLLFANAGMNQFKPIFLNTIDPSHPMAKLSRAANTQKCIRAGGKHNDLDDVGKDVYHHTFFEMLGSWSFGDYFKELACKMALELLTQEFGIPIERLYVTYFGGDEAAGLEADLECKQIWQNLGLDDTKILPGNMKDNFWEMGDTGPCGPCSEIHYDRIGGRDAAHLVNQDDPNVLEIWNLVFIQYNREADGILKPLPKKSIDTGMGLERLVSVLQNKMSNYDTDLFVPYFEAIQKGTGARPYTGKVGAEDADGIDMAYRVLADHARTITVALADGGRPDNTGRGYVLRRILRRAVRYAHEKLNASRGFFATLVDVVVQSLGDAFPELKKDPDMVKDIINEEEVQFLKTLSRGRRILDRKIQSLGDSKTIPGDTAWLLYDTYGFPVDLTGLIAEEKGLVVDMDGFEEERKLAQLKSQGK | P49588 | AARS_M1-K455 | biotinylated recombinant protein |
| alanyl-tRNA synthetase | AlaRS | AARS1 | PL12 | MSA | 1-968 | MDSTLTASEIRQRFIDFFKRNEHTYVHSSATIPLDDPTLLFANAGMNQFKPIFLNTIDPSHPMAKLSRAANTQKCIRAGGKHNDLDDVGKDVYHHTFFEMLGSWSFGDYFKELACKMALELLTQEFGIPIERLYVTYFGGDEAAGLEADLECKQIWQNLGLDDTKILPGNMKDNFWEMGDTGPCGPCSEIHYDRIGGRDAAHLVNQDDPNVLEIWNLVFIQYNREADGILKPLPKKSIDTGMGLERLVSVLQNKMSNYDTDLFVPYFEAIQKGTGARPYTGKVGAEDADGIDMAYRVLADHARTITVALADGGRPDNTGRGYVLRRILRRAVRYAHEKLNASRGFFATLVDVVVQSLGDAFPELKKDPDMVKDIINEEEVQFLKTLSRGRRILDRKIQSLGDSKTIPGDTAWLLYDTYGFPVDLTGLIAEEKGLVVDMDGFEEERKLAQLKSQGKGAGGEDLIMLDIYAIEELRARGLEVTDDSPKYNYHLDSSGSYVFENTVATVMALRREKMFVEEVSTGQECGVVLDKTCFYAEQGGQIYDEGYLVKVDDSSEDKTEFTVKNAQVRGGYVLHIGTIYGDLKVGDQVWLFIDEPRRRPIMSNHTATHILNFALRSVLGEADQKGSLVAPDRLRFDFTAKGAMSTQQIKKAEEIANEMIEAAKAVYTQDCPLAAAKAIQGLRAVFDETYPDPVRVVSIGVPVSELLDDPSGPAGSLTSVEFCGGTHLRNSSHAGAFVIVTEEAIAKGIRRIVAVTGAEAQKALRKAESLKKCLSVMEAKVKAQTAPNKDVQREIADLGEALATAVIPQWQKDELRETLKSLKKVMDDLDRASKADVQKRVLEKTKQFIDSNPNQPLVILEMESGASAKALNEALKLFKMHSPQTSAMLFTVDNEAGKITCLCQVPQNAANRGLKASEWVQQVSGLMDGKGGGKDVSAQATGKNVGCLQEALQLATSFAQLRLGDVKN | P49588 | AARS_M1-N968 | biotinylated recombinant protein |
| alanyl-tRNA synthetase | AlaRS | AARS1 | PL12 | MSA | 830-935 | DRASKADVQKRVLEKTKQFIDSNPNQPLVILEMESGASAKALNEALKLFKMHSPQTSAMLFTVDNEAGKITCLCQVPQNAANRGLKASEWVQQVSGLMDGKGGGKD | P49588 | AARS_HPRR3160056 | PrEST |
| alanyl-tRNA synthetase | AlaRS | AARS1 | PL12 | MSA | 560-672 | EFTVKNAQVRGGYVLHIGTIYGDLKVGDQVWLFIDEPRRRPIMSNHTATHILNFALRSVLGEADQKGSLVAPDRLRFDFTAKGAMSTQQIKKAEEIANEMIEAAKAVYTQDCP | P49588 | AARS_HPRR3160058 | PrEST |
| aminoacyl tRNA synthetase complex interacting multifunctional protein 1 | | AIMP1 | | | 1-312 | MANNDAVLKRLEQKGAEADQIIEYLKQQVSLLKEKAILQATLREEKKLRVENAKLKKEIEELKQELIQAEIQNGVKQIPFPSGTPLHANSMVSENVIQSTAVTTVSSGTKEQIKGGTGDEKKAKEKIEKKGEKKEKKQQSIAGSADSKPIDVSRLDLRIGCIITARKHPDADSLYVEEVDVGEIAPRTVVSGLVNHVPLEQMQNRMVILLCNLKPAKMRGVLSQAMVMCASSPEKIEILAPPNGSVPGDRITFDAFPGEPDKELNPKKKIWEQIQPDLHTNDECVATYKGVPFEVKGKGVCRAQTMSNSGIK | Q12904 | AIMP1_M1-K312 | biotinylated recombinant protein |
| aminoacyl tRNA synthetase complex interacting multifunctional protein 1 | | AIMP1 | | | 211-311 | CNLKPAKMRGVLSQAMVMCASSPEKIEILAPPNGSVPGDRITFDAFPGEPDKELNPKKKIWEQIQPDLHTNDECVATYKGVPFEVKGKGVCRAQTMSNSGI | Q12904 | AIMP1_HPRR2310067 | PrEST |
| aminoacyl tRNA synthetase complex interacting multifunctional protein 1 | | AIMP1 | | | 1-75 | MANNDAVLKRLEQKGAEADQIIEYLKQQVSLLKEKAILQATLREEKKLRVENAKLKKEIEELKQELIQAEIQNGV | Q12904 | AIMP1_HPRR2310068 | PrEST |
| aminoacyl tRNA synthetase complex interacting multifunctional protein 1 | | AIMP1 | | | 84-175 | TPLHANSMVSENVIQSTAVTTVSSGTKEQIKGGTGDEKKAKEKIEKKGEKKEKKQQSIAGSADSKPIDVSRLDLRIGCIITARKHPDADSLY | Q12904 | AIMP1_HPRR2310069 | PrEST |
| aminoacyl tRNA synthetase complex interacting multifunctional protein 1 | | AIMP1 | | | 1-58 | MANNDAVLKRLEQKGAEADQIIEYLKQQVSLLKEKAILQATLREEKKLRVENAKLKKE | Q12904 | AIMP1_HPRR3970191 | PrEST |
| aminoacyl tRNA synthetase complex interacting multifunctional protein 1 | | AIMP1 | | | 147-312 | SKPIDVSRLDLRIGCIITARKHPDADSLYVEEVDVGEIAPRTVVSGLVNHVPLEQMQNRMVILLCNLKPAKMRGVLSQAMVMCASSPEKIEILAPPNGSVPGDRITFDAFPGEPDKELNPKKKIWEQIQPDLHTNDECVATYKGVPFEVKGKGVCRAQTMSNSGIK | Q12904 | AIMP1_S147-K312 | biotinylated recombinant protein |
| aminoacyl tRNA synthetase complex interacting multifunctional protein 2 | | AIMP2 | | | 118-320 | ALKDIVINANPASPPLSLLVLHRLLCEHFRVLSTVHTHSSVKSVPENLLKCFGEQNKKQPRQDYQLGFTLIWKNVPKTQMKFSIQTMCPIEGEGNIARFLFSLFGQKHNAVNATLIDSWVDIAIFQLKEGSSKEKAAVFRSMNSALGKSPWLAGNELTVADVVLWSVLQQIGGCSVTVPANVQRWMRSCENLAPFNTALKLLK | Q13155 | AIMP2_A118-K320 | biotinylated recombinant protein* |
| aminoacyl tRNA synthetase complex interacting multifunctional protein 2 | | AIMP2 | | | 14-106 | APLRVELPTCMYRLPNVHGRSYGPAPGAGHVQEESNLSLQALESRQDDILKRLYELKAAVDGLSKMIQTPDADLDVTNIIQADEPTTLTTNAL | Q13155 | AIMP2_HPRR2440262 | PrEST |
| aminoacyl tRNA synthetase complex interacting multifunctional protein 2 | | AIMP2 | | | 110-196 | SVLGKDYGALKDIVINANPASPPLSLLVLHRLLCEHFRVLSTVHTHSSVKSVPENLLKCFGEQNKKQPRQDYQLGFTLIWKNVPKTQ | Q13155 | AIMP2_HPRR2440263 | PrEST* |
| aminoacyl tRNA synthetase complex interacting multifunctional protein 2 | | AIMP2 | | | 213-320 | IARFLFSLFGQKHNAVNATLIDSWVDIAIFQLKEGSSKEKAAVFRSMNSALGKSPWLAGNELTVADVVLWSVLQQIGGCSVTVPANVQRWMRSCENLAPFNTALKLLK | Q13155 | AIMP2_HPRR2440264 | PrEST* |
| aminoacyl tRNA synthetase complex interacting multifunctional protein 2 | | AIMP2 | | | 90-320 | TNIIQADEPTTLTTNALDLNSVLGKDYGALKDIVINANPASPPLSLLVLHRLLCEHFRVLSTVHTHSSVKSVPENLLKCFGEQNKKQPRQDYQLGFTLIWKNVPKTQMKFSIQTMCPIEGEGNIARFLFSLFGQKHNAVNATLIDSWVDIAIFQLKEGSSKEKAAVFRSMNSALGKSPWLAGNELTVADVVLWSVLQQIGGCSVTVPANVQRWMRSCENLAPFNTALKLLK | Q13155 | AIMP2_T90-K320 | biotinylated recombinant protein* |
| aminoacyl tRNA synthetase complex interacting multifunctional protein 3 | | AIMP3 | | | 7-98 | LSLLEKSLGLSKGNKYSAQGERQIPVLQTNNGPSLTGLTTIAAHLVKQANKEYLLGSTAEEKAIVQQWLEYRVTQVDGHSSKNDIHTLLKDL | O43324 | EEF1E1_HPRR2620052 | PrEST |
| aminoacyl tRNA synthetase complex interacting multifunctional protein 3 | | AIMP3 | | | 123-174 | GLHRFIVDLTVQEKEKYLNVSRWFCHIQHYPGIRQHLSSVVFIKNRLYTNSH | O43324 | EEF1E1_HPRR3890428 | PrEST |
| cysteinyl-tRNA synthetase | CysRS | CARS1 | | | 523-658 | SQCNLYMAARKAVRKRPNQALLENIALYLTHMLKIFGAVEEDSSLGFPVGGPGTSLSLEATVMPYLQVLSEFREGVRKIAREQKVPEILQLSDALRDNILPELGVRFEDHEGLPTVVKLVDRNTLLKEREEKRRVE | P49589 | CARS_HPRR400045 | PrEST |
| cysteinyl-tRNA synthetase | CysRS | CARS1 | | | 110-238 | HLFEQYREKRPEAAQLLEDVQAALKPFSVKLNETTDPDKKQMLERIQHAVQLATEPLEKAVQSRLTGEEVNSCVEVLLEEAKDLLSDWLDSTLGCDVTDNSIFSKLPKFWEGDFHRDMEALNVLPPDVL | P49589 | CARS_HPRR400046 | PrEST |

| | | | | | | | | | |
|---|---|---|---|---|---|---|---|---|---|
| cysteinyl-tRNA synthetase | CysRS | CARS1 | | | 1-748 | MADSSGQQGKGRRVQPQWSPPAGTQPCRLHLYNSLTRNKEVFIPQDGKKVTWYCCGPTVYDASHMGHARSYISFDILRRVLKDYFKFDVFYCMNITDIDDKIIKRARQNHLFEQYREKRPEAAQLLEDVQAALKPFSVKLNETTDPDKKQMLERIQHAVQLATEPLEKAVQSRLTGEEVNSCVEVLLEEAKDLLSDWLDSTLGCDVTDNSIFSKLPKFWEGDFHRDMEALNVLPPDVLTRVSEYVPEIVNFVQKIVDNGYGYVSNGSVYFDTAKFASSEKHSYGKLVPEAVGDQKALQEGEGDLSISADRLSEKRSPNDFALWKASKPGEPSWPCPWGKGRPGWHIECSAMAGTLLGASMDIHGGGFDLRFPHHDNELAQSEAYFENDCWVRYFLHTGHLTIAGCKMSKSLKNFITIKDALKKHSARQLRLAFLMHSWKDTLDYSSNTMESALQYEKFLNEFFLNVKDILRAPVDITGQFEKWGEEEAELNKNFYDKKTAIHKALCDNVDTRTVMEEMRALVSQCNLYMAARKAVRKRPNQALLENIALYLTHMLKIFGAVEEDSSLGFPVGGPGTSLSLEATVMPYLQVLSEFREGVRKIAREQKVPEILQLSDALRDNILPELGVRFEDHEGLPTVVKLVDRNTLLKEREEKRRVEEEKRKKKEEAARRKQEQEAAKLAKMKIPPSEMFLSETDKYSKFDENGLPTHDMEGKELSKGQAKKLKKLFEAQEKLYKEYLQMAQNGSFQ | P49589 | CysRS_M1-Q748 | biotinylated recombinant protein |
| chromodomain helicase DNA binding protein 3 | | CHD3 | Mi-2a | MSA | 292-364 | LKIKLGLLGGKREKGGSYVFQSDEGPEPEAEESDLDSGSVHSASGRPDGPVRTKKLKRGRPGRKKKKLLGCPA | Q12873 | CHD3_HPRR3420199 | PrEST |
| chromodomain helicase DNA binding protein 3 | | CHD3 | Mi-2a | MSA | 1617-1717 | PAPSEKGEGIRTPLEKEEAENQEEKPEKNSRIGEKMETEADAPSPAPSLGERLEPRKIPLEDEVPGVPGEMEPEPGYRGDREKSATESTPGERGEEKPLDG | Q12873 | CHD3_HPRR4290528 | PrEST |
| chromodomain helicase DNA binding protein 4 | | CHD4 | Mi-2b | MSA | 1508-1646 | EFEHVNGRWSMPELAEVEENKKMSQPGSPSPKTPTPSTPGDTQPNTPAPVPPAEDGIKIEENSLKEEESIEGEKEVKSTAPETAIECTQAPAPASEDEKVVVEPPEGEEKVEKAEVKERTEEPMETEPKGAADVEKVEE | Q14839 | CHD4_HPRR680141 | PrEST |
| chromodomain helicase DNA binding protein 4 | | CHD4 | Mi-2b | MSA | 21-141 | DALLNNSLPPPHPENEEDPEEDLSETETPKLKKKKKPKKPRDPKIPKSKRQKKELGDSSGEGPEFVEEEEEVALRSDSEGSDYTPGKKKKKKLGPKKEKKSKSKRKEEEEEDDDDDDSKEP | Q14839 | CHD4_HPRR680142 | PrEST |
| aspartyl-tRNA synthetase | AspRS | DARS1 | | | 392-495 | DKYPLAVRPFYTMPDPRNPKQSNSYDMFMRGEEILSGAQRIHDPQLLTERALHHGIDLEKIKAYIDSFRFGAPPHAGGGIGLERVTMLFLGLHNVRQTSMFPRD | P14868 | DARS_HPRR2370049 | PrEST* |
| aspartyl-tRNA synthetase | AspRS | DARS1 | | | 1-135 | MPSASASRKSQEKPREIMDAAEDYAKERYGISSMIQSQEKPDRVLVRVRDLTIQKADEVVWVRARVHTSRAKGKQCFLVLRQQQFNVQALVAVGDHASKQMVKFAANINKESIVDVEGVVRKVNQKIGSCTQQDV | P14868 | DARS_HPRR2370050 | PrEST |
| aspartyl-tRNA synthetase | AspRS | DARS1 | | | 137-261 | LHVQKIYVISLAEPRLPLQLDDAVRPEAEGEEEGRATVNQDTRLDNRVIDLRTSTSQAVFRLQSGICHLFRETLINKGFVEIQTPKIISAASEGGANVFTVSYFKNNAYLAQSPQLYKQMCICAD | P14868 | DARS_HPRR2370051 | PrEST |
| aspartyl-tRNA synthetase | AspRS | DARS1 | | | 274-385 | AEDSNTHRHLTEFVGLDIEMAFNYHYHEVMEEIADTMVQIFKGLQERFQTEIQTVNKQFPCEPFKFLEPTLRLEYCEALAMLREAGVEMGDEDDLSTPNEKLLGHLVKEKYD | P14868 | DARS_HPRR2370052 | PrEST |
| aspartyl-tRNA synthetase | AspRS | DARS1 | | | 1-501 | MPSASASRKSQEKPREIMDAAEDYAKERYGISSMIQSQEKPDRVLVRVRDLTIQKADEVVWVRARVHTSRAKGKQCFLVLRQQQFNVQALVAVGDHASKQMVKFAANINKESIVDVEGVVRKVNQKIGSCTQQDVELHVQKIYVISLAEPRLPLQLDDAVRPEAEGEEEGRATVNQDTRLDNRVIDLRTSTSQAVFRLQSGICHLFRETLINKGFVEIQTPKIISAASEGGANVFTVSYFKNNAYLAQSPQLYKQMCICADFEKVFSIGPVFRAEDSNTHRHLTEFVGLDIEMAFNYHYHEVMEEIADTMVQIFKGLQERFQTEIQTVNKQFPCEPFKFLEPTLRLEYCEALAMLREAGVEMGDEDDLSTPNEKLLGHLVKEKYDTDFYILDKYPLAVRPFYTMPDPRNPKQSNSYDMFMRGEEILSGAQRIHDPQLLTERALHHGIDLEKIKAYIDSFRFGAPPHAGGGIGLERVTMLFLGLHNVRQTSMFPRDPKRLTP | P14868 | DARS_M1-P501 | biotinylated recombinant protein* |
| glutamatyl-prolyl-tRNA synthetase | EPRS | EPRS1 | | | 1-196 | MATLSLTVNSGDPPLGALLAVEHVKDDVSISVEEGKENILHVSENVIFTDVNSILRYLARVATTAGLYGSNLMEHTEIDHWLEFSATKLSSCDSFTSTINELNHCLSLRTYLVGNSLSLADLCVWATLKGNAAWQEQLKQKKAPVHVKRWFGFLEAQQAFQSVGTKWDVSTTKARVAPEKKQDVGKFVELPGAEMG | P07814 | EPRS_M1-G196 | biotinylated recombinant protein* |
| glutamatyl-prolyl-tRNA synthetase | EPRS | EPRS1 | | | 1026-1113 | DWYSQVITKSEMIEYHDISGCYILRPWAYAIWEAIKDFFDAEIKKLGVENCYFPMFVSQSALEKEKTHVADFAPEVAWVTRSGKTELA | P07814 | EPRS_HPRR2550837 | PrEST* |
| glutamatyl-prolyl-tRNA synthetase | EPRS | EPRS1 | | | 1197-1298 | YEELLAIPVVKGRKTEKEKFAGGDYTTTIEAFISASGRAIQGGTSHHLGQNFSKMFEIVFEDPKIPGEKQFAYQNSWGLTTRTIGVMTMVHGDNMGLVLPPR | P07814 | EPRS_HPRR2550838 | PrEST |
| glutamatyl-prolyl-tRNA synthetase | EPRS | EPRS1 | | | 1301-1405 | CVQVVIIPCGITNALSEEDKEALIAKCNDYRRRLLSVNIRVRADLRDNYSPGWKFNHWELKGVPIRLEVGPRDMKSCQFVAVRRDTGEKLTVAENEAETKLQAIL | P07814 | EPRS_HPRR2550839 | PrEST* |
| exosome component 10 | | EXOSC 10 | PM-Scl 100 | MAA | 102-177 | KVTELEDKFDLLVDANDVILERVGILLDEASGVNKNQQPVLPAGLQVPKTVVSSWNRKAAEYGKKAKSETFRLLHA | Q01780 | EXOSC10_HPRR2551653 | PrEST |
| exosome component 10 | | EXOSC 10 | PM-Scl 100 | MAA | 254-338 | FAHPYQYELNHFTPADAVLQKPQPQLYRPIEETPCHFISSLDELVELNEKLLNCQEFAVDLEHHSYRSFLGLTCLMQISTRTEDF | Q01780 | EXOSC10_HPRR2551654 | PrEST |
| exosome component 9 | | EXOSC 9 | PM-Scl 75 | MAA | 122-197 | CVVAGEKVWQIRVDLHLLNHDGNIIDAASIAAIVALCHFRRPDVSVQGDEVTLYTPEERDPVPLSIHHMPICVSFA | Q06265 | EXOSC9_HPRR3000153 | PrEST |
| exosome component 9 | | EXOSC 9 | PM-Scl 75 | MAA | 209-295 | PNEREERVMDGLLVIAMNKHREICTIQSSGGIMLLKDQVLRCSKIAGVKVAEITELILKALENDQKVRKEGGKFGFAESIANQRITA | Q06265 | EXOSC9_HPRR3000155 | PrEST |
| phenylalanyl-tRNA synthetase alpha subunit | PheRSa | FARSA | Zo | MSA | 1-182 | MADGQVAELLLRRLEASDGGLDSAELAAELGMEHQAVVGAVKSLQALGEVIEAELRSTKHWELTAEGEEIAREGSHEARVFRSIPPEGLAQSELMRLPSGKVGFSKAMSNKWIRVDKSAADGPRVFRVVDSMEDEVQRRLQLVRGGQAEKLGEKERSELRKRKLLAEVTLKTYWVSKGSAFS | Q9Y285 | FARSA_M1-S182 | biotinylated recombinant protein |
| phenylalanyl-tRNA synthetase alpha subunit | PheRSa | FARSA | Zo | MSA | 49-178 | EVIEAELRSTKHWELTAEGEEIAREGSHEARVFRSIPPEGLAQSELMRLPSGKVGFSKAMSNKWIRVDKSAADGPRVFRVVDSMEDEVQRRLQLVRGGQAEKLGEKERSELRKRKLLAEVTLKTYWVSKG | Q9Y285 | FARSA_HPRR400146 | PrEST |
| phenylalanyl-tRNA synthetase alpha subunit | PheRSa | FARSA | Zo | MSA | 325-407 | RTHTTSASARALYRLAQKKPFTPVKYFSIDRVFRNETLDATHLAEFHQIEGVVADHGLTLGHLMGVLREFFTKLGITQLRFKP | Q9Y285 | FARSA_HPRR4240045 | PrEST |
| phenylalanyl-tRNA synthetase beta subunit | PheRSb | FARSB | Zo | MSA | 253-336 | IECTGTDFTKAKIVLDIIVTMFSEYCENQFTVEAAEVVFPNGKSHTFPELAYRKEMVRADLINKKVGIRETPENLAKLLTRMYL | Q9NSD9 | FARSB_HPRR2920307 | PrEST* |

| | | | | | | | | | |
|---|---|---|---|---|---|---|---|---|---|
| phenylalanyl-tRNA synthetase beta subunit | PheRSb | FARSB | Zo | MSA | 161-248 | THDLDTLSGPFTYTAKRPSDIKFKPLNKTKEYTACELMNIYKTDNHLKHYLHIIENKPLYPVIYDSNGVVLSMPPIINGDHSRITVNT | Q9NSD9 | FARSB_HPRR2920308 | PrEST* |
| phenylalanyl-tRNA synthetase beta subunit | PheRSb | FARSB | Zo | MSA | 46-146 | SKEQGNVKAAGASDVVLYKIDVPANRYDLLCLEGLVRGLQVFKERIKAPVYKRVMPDGKIQKLIITEETAKIRPFAVAAVLRNIKFTKDRYDSFIELQEKL | Q9NSD9 | FARSB_HPRR2920309 | PrEST |
| phenylalanyl-tRNA synthetase subunit beta | PheRSb | FARSB | Zo | MSA | 1-589 | MPTVSVKRDLLFQALGRTYTDEEFDELCFEFGLELDEITSEKEIISKEQGNVKAAGASDVVLYKIDVPANRYDLLCLEGLVRGLQVFKERIKAPVYKRVMPDGKIQKLIITEETAKIRPFAVAAVLRNIKFTKDRYDSFIELQEKLHQNICRKRALVAIGTHDLDTLSGPFTYTAKRPSDIKFKPLNKTKEYTACELMNIYKTDNHLKHYLHIIENKPLYPVIYDSNGVVLSMPPIINGDHSRITVNTRNIFIECTGTDFTKAKIVLDIIVTMFSEYCENQFTVEAAEVVFPNGKSHTFPELAYRKEMVRADLINKKVGIRETPENLAKLLTRMYLKSEVIGDGNQIEIEIPPTRADIIHACDIVEDAAIAYGYNNIQMTLPKTYTIANQFPLNKLTELLRHDMAAAGFTEALTFALCSQEDIADKLGVDISATKAVHISNPKTAEFQVARTTLLPGLLKTIAANRKMPLPLKLFEISDIVIKDSNTDVGAKNYRHLCAVYYNKNPGFEIIHGLLDRIMQLLDVPPGEDKGGYVIKASEGPAFFPGRCAEIFARGQSVGKLGVLHPDVITKFELTMPCSSLEINIGPFL | Q9NSD9 | FARSB_M1-L589 | biotinylated recombinant protein |
| glycyl-tRNA synthetase | GlyRS | GARS1 | EJ | MSA | 56-117 | DGAGAEEVLAPLRLAVRQQGDLVRKLKEDKAPQVDVDKAVAELKARKRVLEAKELALQPKDD | P41250 | GARS_D56-D117 | biotinylated recombinant protein |
| glycyl-tRNA synthetase | GlyRS | GARS1 | EJ | MSA | 224-326 | KLMSDKKCSVEKKSEMESVLAQLDNYGQQELADLFVNYNVKSPITGNDLSPPVSFNLMFKTFIGPGGNMPGYLRPETAQGIFLNFKRLLEFNQGKLPFAAAQI | P41250 | GARS_HPRR2440240 | PrEST* |
| glycyl-tRNA synthetase | GlyRS | GARS1 | EJ | MSA | 619-728 | LPLSQNQEFMPFVKELSEALTRHGVSHKVDDSSGSIGRRYARTDEIGVAFGVTIDFDTVNKTPHTATLRDRDSMRQIRAEISELPSIVQDLANGNITWADVEARYPLFEG | P41250 | GARS_HPRR2440243 | PrEST |
| glycyl-tRNA synthetase | GlyRS | GARS1 | EJ | MSA | 55-739 | DGAGAEEVLAPLRLAVRQQGDLVRKLKEDKAPQVDVDKAVAELKARKRVLEAKELALQPKDDIVDRAKMEDTLKRRFFYDQAFAIYGGVSGLYDFGPVGCALKNNIIQTWRQHFIQEEQILEIDCTMLTPEPVLKTSGHVDKFADFMVKDVKNGECFRADHLLKAHLQKLMSDKKCSVEKKSEMESVLAQLDNYGQQELADLFVNYNVKSPITGNDLSPPVSFNLMFKTFIGPGGNMPGYLRPETAQGIFLNFKRLLEFNQGKLPFAAAQIGNSFRNEISPRSGLIRVREFTMAEIEHFVDPSEKDHPKFQNVADLHLYLYSAKAQVSGQSARKMRLGDAVEQGVINNTVLGYFIGRIYLYLTKVGISPDKLRFRQHMENEMAHYACDCWDAESKTSYGWIEIVGCADRSCYDLSCHARATKVPLVAEKPLKEPKTVNVVQFEPSKGAIGKAYKKDAKLVMEYLAICDECYITEMEMLLNEKGEFTIETEGKTFQLTKDMINVKRFQKTLYVEEVVPNVIEPSFGLGRIMYTVFEHTFHVREGDEQRTFFSFPAVVAPFKCSVLPLSQNQEFMPFVKELSEALTRHGVSHKVDDSSGSIGRRYARTDEIGVAFGVTIDFDTVNKTPHTATLRDRDSMRQIRAEISELPSIVQDLANGNITWADVEARYPLFEGQETGKKETIEE | P41250 | GARS_M55-E739 | biotinylated recombinant protein |
| histidyl-tRNA synthetase | HisRS | HARS1 | Jo1 | MSA | 1-509 | MAERAALEELVKLQGERVRGLKQQKASAELIEEEVAKLLKLKAQLGPDESKQKFVLKTPKGTRDYSPRQMAVREKVFDVIIRCFKRHGAEVIDTPVFELKETLMGKYGEDSKLIYDLKDQGGELLSLRYDLTVPFARYLAMNKLTNIKRYHIAKVYRRDNPAMTRGRYREFYQCDFDIAGNFDPMIPDAECLKIMCEILSSLQIGDFLVKVNDRRILDGMFAICGVSDSKFRTICSSVDKLDKVSWEEVKNEMVGEKGLAPEVADRIGDYVQQHGGVSLVEQLLQDPKLSQNKQALEGLGDLKLLFEYLTLFGIDDKISFDLSLARGLDYYTGVIYEAVLLQTPAQAGEEPLGVGSVAAGGRYDGLVGMFDPKGRKVPCVGLSIGVERIFSIVEQRLEALEEKIRTTETQVLVASAQKKLLEERLKLVSELWDAGIKAELLYKKNPKLLNQLQYCEEAGIPLVAIIGEQELKDGVIKLRSVTSREEVDVRREDLVEEIKRRTGQPLCIC | P12081 | HARS_M1-C509 | biotinylated recombinant protein |
| histidyl-tRNA synthetase | HisRS | HARS1 | Jo1 | MSA | 1-60 | MAERAALEELVKLQGERVRGLKQQKASAELIEEEVAKLLKLKAQLGPDESKQKFVLKTPK | P12081 | HARS_M1-K60 | biotinylated recombinant protein |
| histidyl-tRNA synthetase | HisRS | HARS1 | Jo1 | MSA | 1-64 | MAERAALEELVKLQGERVRGLKQQKASAELIEEEVAKLLKLKAQLGPDESKQKFVLKTPKGTRD | P12081 | HARS_HPRR3010485 | PrEST |
| histidyl-tRNA synthetase | HisRS | HARS1 | Jo1 | MSA | 473-509 | DGVIKLRSVTSREEVDVRREDLVEEIKRRTGQPLCIC | P12081 | HARS_HPRR4040036 | PrEST |
| histidyl-tRNA synthetase | HisRS | HARS1 | Jo1 | MSA | 53-509 | KFVLKTPKGTRDYSPRQMAVREKVFDVIIRCFKRHGAEVIDTPVFELKETLMGKYGEDSKLIYDLKDQGGELLSLRYDLTVPFARYLAMNKLTNIKRYHIAKVYRRDNPAMTRGRYREFYQCDFDIAGNFDPMIPDAECLKIMCEILSSLQIGDFLVKVNDRRILDGMFAICGVSDSKFRTICSSVDKLDKVSWEEVKNEMVGEKGLAPEVADRIGDYVQQHGGVSLVEQLLQDPKLSQNKQALEGLGDLKLLFEYLTLFGIDDKISFDLSLARGLDYYTGVIYEAVLLQTPAQAGEEPLGVGSVAAGGRYDGLVGMFDPKGRKVPCVGLSIGVERIFSIVEQRLEALEEKIRTTETQVLVASAQKKLLEERLKLVSELWDAGIKAELLYKKNPKLLNQLQYCEEAGIPLVAIIGEQELKDGVIKLRSVTSREEVDVRREDLVEEIKRRTGQPLCIC | P12081 | HARS_K53-C509 | biotinylated recombinant protein |
| histidyl-tRNA synthetase | HisRS | HARS1 | Jo1 | MSA | 53-402 | KGTRDYSPRQMAVREKVFDVIIRCFKRHGAEVIDTPVFELKETLMGKYGEDSKLIYDLKDQGGELLSLRYDLTVPFARYLAMNKLTNIKRYHIAKVYRRDNPAMTRGRYREFYQCDFDIAGNFDPMIPDAECLKIMCEILSSLQIGDFLVKVNDRRILDGMFAICGVSDSKFRTICSSVDKLDKVSWEEVKNEMVGEKGLAPEVADRIGDYVQQHGGVSLVEQLLQDPKLSQNKQALEGLGDLKLLFEYLTLFGIDDKISFDLSLARGLDYYTGVIYEAVLLQTPAQAGEEPLGVGSVAAGGRYDGLVGMFDPKGRKVPCVGLSIGVERIFSIVEQRLEALEE | P12081 | HARS_K53-E402 | biotinylated recombinant protein |
| histidyl-tRNA synthetase | HisRS | HARS1 | Jo1 | MSA | 1-60_399-509 | MAERAALEELVKLQGERVRGLKQQKASAELIEEEVAKLLKLKAQLGPDESKQKFVLKTPKALEEKIRTTETQVLVASAQKKLLEERLKLVSELWDAGIKAELLYKKNPKLLNQLQYCEEAGIPLVAIIGEQELKDGVIKLRSVTSREEVDVRREDLVEEIKRRTGQPLCIC | P12081 | HARS_M1-K60_A399-C509 | biotinylated recombinant protein |
| histidyl-tRNA synthetase | HisRS | HARS1 | Jo1 | MSA | 406-509 | TTETQVLVASAQKKLLEERLKLVSELWDAGIKAELLYKKNPKLLNQLQYCEEAGIPLVAIIGEQELKDGVIKLRSVTSREEVDVRREDLVEEIKRRTGQPLCIC | P12081 | HARS_T406-C509 | biotinylated recombinant protein |
| isoleucyl-tRNA synthetase | IleRS | IARS1 | OJ | MSA | 732-831 | NWYVRMNRRRLKGENGMEDCVMALETLFSVLLSLCRLMAPYTPFLTELMYQNLKVLIDPVSVQDKDTLSIHYLMLPRVREELIDKKTESAVSQMQSVIEL | P41252 | IARS_HPRR2470884 | PrEST |
| isoleucyl-tRNA synthetase | IleRS | IARS1 | OJ | MSA | 987-1099 | AREVINRIQKLRKKCNLVPTDEITVYYKAKSEGTYLNSVIESHTEFIFTTIKAPLKPYPVSPSDKVLIQEKTQLKGSELEITLTRGSSLPGPACAYVNLNICANGSEQGGVLL | P41252 | IARS_HPRR2470886 | PrEST |
| lysyl-tRNA synthetase | LysRS | KARS1 | | | 1-597 | MAAVQAAEVKVDGSEPKLSKNELKRRLKAEKKVAEKEAKQKELSEKQLSQATAAATNHTTDNGVGPEEESVDPNQYYKIRSQAIHQLKVNGEDPYPHKFHVDISLTDFIQKYSHLQPGDHLTDITLKVAGRIHAKRASGGKLIFYDLRGEGVKLQVMANSRNYKSEEEFIHINNKLRRGDIIGVQGNPGKTKKGELSIIPYEITLLSPCLHMLPHLHFGLKDKETRYRQRYLDLILNDFVRQKFIIRSKIITYIRSFLDELGFLEIETPMMNIIPGGAVAKPFITYHNELDMNLYMRIAPELYHKMLVVGGIDRVYEIGRQFRNEGIDLTHNPEFTTCEFYMAYADYHDLMEITEKMVSGMVKHITGSYKVTYHPDGPEGQAYDVDFTPPFRRINMVEELEKALGMKLPETNLFETEETRKILDDICVAKAVECPPPRTTARLLDKLVGEFLEVTCINPTFICDHPQIMSPLAKWHRSKEGLTERFELFVMKKEICNAYTELNDPMRQRQLFEEQAKAKAAGDDEAMFIDENFCTALEYGLPPTAGWGMGIDRVAMFLTDSNNIKEVLLFPAMKPEDKKENVATTDTLESTTVGTSV | Q15046 | KARS_M1-V597 | biotinylated recombinant protein |

| | | | | | | | | | |
|---|---|---|---|---|---|---|---|---|---|
| lysyl-tRNA synthetase | LysRS | KARS1 | | | 360-479 | GMVKHITGSYKVTYHPDGPEGQAYDVDFTPPFRRINMVEELEKALGMKLPETNLFETEETRKILDDICVAKAVECPPPRTTARLLDKLVGEFLEVTCINPTFICDHPQIMSPLAKWHRSK | Q15046 | KARS_HPRR3150005 | PrEST* |
| lysyl-tRNA synthetase | LysRS | KARS1 | | | 481-591 | GLTERFELFVMKKEICNAYTELNDPMRQRQLFEEQAKAKAAGDDEAMFIDENFCTALEYGLPPTAGWGMGIDRVAMFLTDSNNIKEVLLFPAMKPEDKKENVATTDTLEST | Q15046 | KARS_HPRR3150006 | PrEST |
| leucyl-tRNA synthetase | LeuRS | LARS1 | | | 1-1176 | MAERKGTAKVDFLKKIEKEIQQKWDTERVFEVNASNLEKQTSKGKYFVTFPYPYMNGRLHLGHTFSLSKCEFAVGYQRLKGKCCLFPFGLHCTGMPIKACADKLKREIELYGCPPDFPDEEEEEEETSVKTEDIIIKDKAKGKKSKAAAKAGSSKYQWGIMKSLGLSDEEIVKFSEAEHWLDYFPPLAIQDLKRMGLKVDWRRSFITTDVNPYYDSFVRWQFLTLRERNKIKFGKRYTIYSPKDGQPCMDHDRQTGEGVGPQEYTLLKLKVLEPYPSKLSGLKGKNIFLVAATLRPETMFGQTNCWVRPDMKYIGFETVNGDIFICTQKAARNMSYQGFTKDNGVVPVVKELMGEEILGASLSAPLTSYKVIYVLPMLTIKEDKGTGVVTSVPSDSPDDIAALRDLKKKQALRAKYGIRDDMVLPFEPVPVIEIPGFGNLSAVTICDELKIQSQNDREKLAEAKEKIYLKGFYEGIMLVDGFKGQKVQDVKKTIQKKMIDAGDALIYMEPEKQVMSRSSDECVVALCDQWYLDYGEENWKKQTSQCLKNLETFCEETRRNFEATLGWLQEHACSRTYGLGTHLPWDEQWLIESLSDSTIYMAFYTVAHLLQGGNLHGQAESPLGIRPQQMTKEVWDYVFFKEAPFPKTQIAKEKLDQLKQEFEFWYPVDLRVSGKDLVPNHLSYYLYNHVAMWPEQSDKWPTAVRANGHLLLNSEKMSKSTGNFLTLTQAIDKFSADGMRLALADAGDTVEDANFVEAMADAGILRLYTWVEWVKEMVANWDSLRSGPASTFNDRVFASELNAGIIKTDQNYEKMMFKEALKTGFFEFQAAKDKYRELAVEGMHRELVFRFIEVQTLLLAPFCPHLCEHIWTLLGKPDSIMNASWPVAGPVNEVLIHSSQYLMEVTHDLRLRLKNYMMPAKGKKTDKQPLQKPSHCTIYVAKNYPPWQHTTLSVLRKHFEANNGKLPDNKVIASELGSMPELKKYMKKVMPFVAMIKENLEKMGPRILDLQLEFDEKAVLMENIVYLTNSLELEHIEVKFASEAEDKIREDCCPGKPLNVFRIEPGVSVSLVNPQPSNGHFSTKIEIRQGDNCDSIIRRLMKMNRGIKDLSKVKLMRFDDPLLGPRRVPVLGKEYTEKTPISEHAVFNVDLMSKKIHLTENGIRVDIGDTIIYLVH | Q9P2J5 | LARS_M1-H1176 | biotinylated recombinant protein |
| leucyl-tRNA synthetase | LeuRS | LARS1 | | | 260-511 | GPQEYTLLKLKVLEPYPSKLSGLKGKNIFLVAATLRPETMFGQTNCWVRPDMKYIGFETVNGDIFICTQKAARNMSYQGFTKDNGVVPVVKELMGEEILGASLSAPLTSYKVIYVLPMLTIKEDKGTGVVTSVPSDSPDDIAALRDLKKKQALRAKYGIRDDMVLPFEPVPVIEIPGFGNLSAVTICDELKIQSQNDREKLAEAKEKIYLKGFYEGIMLVDGFKGQKVQDVKKTIQKKMIDAGDALIYMEPE | Q9P2J5 | LARS_G260-E511 | biotinylated recombinant protein |
| leucyl-tRNA synthetase | LeuRS | LARS1 | | | 287-382 | IFLVAATLRPETMFGQTNCWVRPDMKYIGFETVNGDIFICTQKAARNMSYQGFTKDNGVVPVVKELMGEEILGASLSAPLTSYKVIYVLPMLTIKE | Q9P2J5 | LARS_HPRR3010272 | PrEST |
| leucyl-tRNA synthetase | LeuRS | LARS1 | | | 178-272 | EHWLDYFPPLAIQDLKRMGLKVDWRRSFITTDVNPYYDSFVRWQFLTLRERNKIKFGKRYTIYSPKDGQPCMDHDRQTGEGVGPQEYTLLKLKVL | Q9P2J5 | LARS_HPRR3010273 | PrEST |
| leucyl-tRNA synthetase | LeuRS | LARS1 | | | 23-105 | KWDTERVFEVNASNLEKQTSKGKYFVTFPYPYMNGRLHLGHTFSLSKCEFAVGYQRLKGKCCLFPFGLHCTGMPIKACADKLK | Q9P2J5 | LARS_HPRR3010274 | PrEST |
| methionyl-tRNA synthetase | MetRS | MARS1 | | | 271-368 | ALPYVNNVPHLGNIIGCVLSADVFARYSRLRQWNTLYLCGTDEYGTATETKALEEGLTPQEICDKYHIIHADIYRWFNISFDIFGRTTTPQQTKITQD | P56192 | MARS_HPRR4220335 | PrEST* |
| methionyl-tRNA synthetase | MetRS | MARS1 | | | 378-520 | FVLQDTVEQLRCEHCARFLADRFVEGVCPFCGYEEARGDQCDKCGKLINAVELKKPQCKVCRSCPVVQSSQHLFLDLPKLEKRLEEWLGRTLPGSDWTPNAQFITRSWLRDGLKPRCITRDLKWGTPVPLEGFEDKVFYVWFD | P56192 | MARS_HPRR430074 | PrEST* |
| methionyl-tRNA synthetase | MetRS | MARS1 | | | 1-225 | MRLFVSDGVPGCLPVLAAAGRARGRAEVLISTVGPEDCVVPFLTRPKVPVLQLDSGNYLFSTSAICRYFFLLSGWEQDDLTNQWLEWEATELQPALSAALYYLVVQGKKGEDVLGSVRRALTHIDHSLSRQNCPFLAGETESLADIVLWGALYPLLQDPAYLPEELSALHSWFQTLSTQEPCQRAAETVLKQQGVLALRPYLQKQPQPSPAEGRAVTNEPEEEEL | P56192 | MARS_M1-L225 | biotinylated recombinant protein |
| interferon-induced helicase C domain-containing protein 1 | MDA5 | IFIH1 | MDA5 | MSA | 110-207 | AHDEYLQLLNLLQPTLVDKLLVRDVLDKCMEEELLTIEDRNRIAAAENNGNESGVRELLKRIVQKENWFSAFLNVLRQTGNNELVQELTGSDCSESNA | Q9BYX4 | MDA5_A110-A207 | biotinylated recombinant protein |
| interferon-induced helicase C domain-containing protein 1 | MDA5 | IFIH1 | MDA5 | MSA | 110-1025 | AHDEYLQLLNLLQPTLVDKLLVRDVLDKCMEEELLTIEDRNRIAAAENNGNESGVRELLKRIVQKENWFSAFLNVLRQTGNNELVQELTGSDCSESNAEIENLSQVDGPQVEEQLLSTTVQPNLEKEVWGMENNSSESSFADSSVVSESDTSLAEGSVSCLDESLGHNSNMGSDSGTMGSDSDEENVAARASPEPELQLRPYQMEVAQPALEGKNIIICLPTGSGKTRVAVYIAKDHLDKKKKASEPGKVIVLVNKVLLVEQLFRKEFQPFLKKWYRVIGLSGDTQLKISFPEVVKSCDIIISTAQILENSLLNLENGEDAGVQLSDFSLIIIDECHHTNKEAVYNNIMRHYLMQKLKNNRLKKENKPVIPLPQILGLTASPGVGGATKQAKAEEHILKLCANLDAFTIKTVKENLDQLKNQIQEPCKKFAIADATREDPFKEKLLEIMTRIQTYCQMSPMSDFGTQPYEQWAIQMEKKAAKEGNRKERVCAEHLRKYNEALQINDTIRMIDAYTHLETFYNEEKDKKFAVIEDDSDEGGDDEYCDGDEDEDDLKKPLKLDETDRFLMTLFFENNKMLKRLAENPEYENEKLTKLRNTIMEQYTRTEESARGIIFTKTRQSAYALSQWITENEKFAEVGVKAHHLIGAGHSSEFKPMTQNEQKEVISKFRTGKINLLIATTVAEEGLDIKECNIVIRYGLVTNEIAMVQARGRARADESTYVLVAHSGSGVIERETVNDFREKMMYKAIHCVQNMKPEEYAHKILELQMQSIMEKKMKTKRNIAKHYKNNPSLITFLCKNCSVLACSGEDIHVIEKMHHVNMTPEFKELYIVRENKTLQKKCADYQINGEIICKCGQAWGTMMVHKGLDLPCLKIRNFVVVFKNNSTKKQYKKWVELPITFPNLDYSECCLFSDED | Q9BYX4 | MDA5_A110-D1025 | biotinylated recombinant protein |
| interferon-induced helicase C domain-containing protein 1 | MDA5 | IFIH1 | MDA5 | MSA | 110-509 | AHDEYLQLLNLLQPTLVDKLLVRDVLDKCMEEELLTIEDRNRIAAAENNGNESGVRELLKRIVQKENWFSAFLNVLRQTGNNELVQELTGSDCSESNAEIENLSQVDGPQVEEQLLSTTVQPNLEKEVWGMENNSSESSFADSSVVSESDTSLAEGSVSCLDESLGHNSNMGSDSGTMGSDSDEENVAARASPEPELQLRPYQMEVAQPALEGKNIIICLPTGSGKTRVAVYIAKDHLDKKKKASEPGKVIVLVNKVLLVEQLFRKEFQPFLKKWYRVIGLSGDTQLKISFPEVVKSCDIIISTAQILENSLLNLENGEDAGVQLSDFSLIIIDECHHTNKEAVYNNIMRHYLMQKLKNNRLKKENKPVIPLPQILGLTASPGVGGATKQAKAEEHILKL | Q9BYX4 | MDA5_A110-L509 | biotinylated recombinant protein |
| interferon-induced helicase C domain-containing protein 1 | MDA5 | IFIH1 | MDA5 | MSA | 298-1025 | ARASPEPELQLRPYQMEVAQPALEGKNIIICLPTGSGKTRVAVYIAKDHLDKKKKASEPGKVIVLVNKVLLVEQLFRKEFQPFLKKWYRVIGLSGDTQLKISFPEVVKSCDIIISTAQILENSLLNLENGEDAGVQLSDFSLIIIDECHHTNKEAVYNNIMRHYLMQKLKNNRLKKENKPVIPLPQILGLTASPGVGGATKQAKAEEHILKLCANLDAFTIKTVKENLDQLKNQIQEPCKKFAIADATREDPFKEKLLEIMTRIQTYCQMSPMSDFGTQPYEQWAIQMEKKAAKEGNRKERVCAEHLRKYNEALQINDTIRMIDAYTHLETFYNEEKDKKFAVIEDDSDEGGDDEYCDGDEDEDDLKKPLKLDETDRFLMTLFFENNKMLKRLAENPEYENEKLTKLRNTIMEQYTRTEESARGIIFTKTRQSAYALSQWITENEKFAEVGVKAHHLIGAGHSSEFKPMTQNEQKEVISKFRTGKINLLIATTVAEEGLDIKECNIVIRYGLVTNEIAMVQARGRARADESTYVLVAHSGSGVIERETVNDFREKMMYKAIHCVQNMKPEEYAHKILELQMQSIMEKKMKTKRNIAKHYKNNPSLITFLCKNCSVLACSGEDIHVIEKMHHVNMTPEFKELYIVRENKTLQKKCADYQINGEIICKCGQAWGTMMVHKGLDLPCLKIRNFVVVFKNNSTKKQYKKWVELPITFPNLDYSECCLFSDED | Q9BYX4 | MDA5_A298-D1025 | biotinylated recombinant protein |
| interferon-induced helicase C domain-containing protein 1 | MDA5 | IFIH1 | MDA5 | MSA | 816-1025 | MVQARGRARADESTYVLVAHSGSGVIERETVNDFREKMMYKAIHCVQNMKPEEYAHKILELQMQSIMEKKMKTKRNIAKHYKNNPSLITFLCKNCSVLACSGEDIHVIEKMHHVNMTPEFKELYIVRENKTLQKKCADYQINGEIICKCGQAWGTMMVHKGLDLPCLKIRNFVVVFKNNSTKKQYKKWVELPITFPNLDYSECCLFSDED | Q9BYX4 | MDA5_M816-D1025 | biotinylated recombinant protein |

| | | | | | | | | | |
|---|---|---|---|---|---|---|---|---|---|
| interferon-induced helicase C domain-containing protein 1 | MDA5 | IFIH1 | MDA5 | MSA | 896-1025 | YKNNPSLITFLCKNCSVLACSGEDIHVIEKMHHVNMTPEFKELYIVRENKTLQKKCADYQINGEIICKCGQAWGTMMVHKGLDLPCLKIRNFVVVFKNNSTKKQYKKWVELPITFPNLDYSECCLFSDED | Q9BYX4 | MDA5_Y896-D1025 | biotinylated recombinant protein |
| interferon-induced helicase C domain-containing protein 1 | MDA5 | IFIH1 | MDA5 | MSA | 74-223 | GWTREFVEALRRTGSPLAARYMNPELTDLPSPSFENAHDEYLQLLNLLQPTLVDKLLVRDVLDKCMEEELLTIEDRNRIAAAENNGNESGVRELLKRIVQKENWFSAFLNVLRQTGNNELVQELTGSDCSESNAEIENLSQVDGPQVEEQ | Q9BYX4 | IFIH1_HPRR330271 | PrEST |
| asparaginyl-tRNA synthetase | AsnRS | NARS1 | KS | MSA | 353-445 | RLEDLVCDVVDRILKSPAGSIVHELNPNFQPPKRPFKRMNYSDAIVWLKEHDVKKEDGTFYEFGEDIPEAPERLMTDTINEPILLCRFPVEIK | O43776 | NARS_HPRR3120288 | PrEST |
| asparaginyl-tRNA synthetase | AsnRS | NARS1 | KS | MSA | 118-209 | GALEGYRGQRVKVFGWVHRLRRQGKNLMFLVLRDGTGYLQCVLADELCQCYNGVLLSTESSVAVYGMLNLTPKGKQAPGGHELSCDFWELIG | O43776 | NARS_HPRR3120291 | PrEST |
| asparaginyl-tRNA synthetase | AsnRS | NARS1 | KS | MSA | 1-548 | MVLAELYVSDREGSDATGDGTKEKPFKTGLKALMTVGKEPFPTIYVDSQKENERWNVISKSQLKNIKKMWHREQMKSESREKKEAEDSLRREKNLEEAKKITIKNDPSLPEPKCVKIGALEGYRGQRVKVFGWVHRLRRQGKNLMFLVLRDGTGYLQCVLADELCQCYNGVLLSTESSVAVYGMLNLTPKGKQAPGGHELSCDFWELIGLAPAGGADNLINEESDVDVQLNNRHMMIRGENMSKILKARSMVTRCFRDHFFDRGYYEVTPPTLVQTQVEGGATLFKLDYFGEEAFLTQSSQLYLETCLPALGDVFCIAQSYRAEQSRTRRHLAEYTHVEAECPFLTFDDLLNRLEDLVCDVVDRILKSPAGSIVHELNPNFQPPKRPFKRMNYSDAIVWLKEHDVKKEDGTFYEFGEDIPEAPERLMTDTINEPILLCRFPVEIKSFYMQRCPEDSRLTESVDVLMPNVGEIVGGSMRIFDSEEILAGYKREGIDPTPYYWYTDQRKYGTCPHGGYGLGLERFLTWILNRYHIRDVCLYPRFVQRCTP | O43776 | NARS_M1-P548 | biotinylated recombinant protein |
| glutaminyl-tRNA synthetase | GlnRS | QARS1 | | | A2-V775 | AALDSLSLFTSLGLSEQKARETLKNSALSAQLREAATQAQQTLGSTIDKATGILLYGLASRLRDTRRLSFLVSYIASKKIHTEPQLSAALEYVRSHPLDPIDTVDFERECGVGVIVTPEQIEEAVEAAINRHRPQLLVERYHFNMGLLMGEARAVLKWADGKMIKNEVDMQVLHLLGPKLEADLEKKFKVAKARLEETDRRTAKDVVENGETADQTLSLMEQLRGEALKFHKPGENYKTPGYVVTPHTMNLLKQHLEITGGQVRTRFPPEPNGILHIGHAKAINFNFGYAKANNGICFLRFDDTNPEKEEAKFFTAICDMVAWLGYTPYKVTYASDYFDQLYAWAVELIRRGLAYVCHQRGEELKGHNTLPSPWRDRPMEESLLLFEAMRKGKFSEGEATLRMKLVMEDGKMDPVAYRVKYTPHHRTGDKWCIYPTYDYTHCLCDSIEHITHSLCTKEFQARRSSYFWLCNALDVYCPVQWEYGRLNLHYAVVSKRKILQLVATGAVRDWDDPRLFTLTALRRRGFPPEAINNFCARVGVTVAQTTMEPHLLEACVRDVLNDTAPRAMAVLESLRVIITNFPAAKSLDIQVPNFPADETKGFHQVPFAPIVFIERTDFKEEPEPGFKRLAWGQPVGLRHTGYVIELQHVVKGPSGCVESLEVTCRRADAGEKPKAFIHWVSQPLMCEVRLYERLFQHKNPEDPTEVPGGFLSDLNLASLHVVDAALVDCSVALAKPFDKFQFERLGYFSVDPDSHQGKLVFNRTVTLKEDPGKV | P47897 | QARS_M1-V775 | biotinylated recombinant protein |
| glutaminyl-tRNA synthetase | GlnRS | QARS1 | | | 640-725 | HTGYVIELQHVVKGPSGCVESLEVTCRRADAGEKPKAFIHWVSQPLMCEVRLYERLFQHKNPEDPTEVPGGFLSDLNLASLHVVDA | P47897 | QARS_HPRR2960726 | PrEST |
| glutaminyl-tRNA synthetase | GlnRS | QARS1 | | | 69-169 | LSFLVSYIASKKIHTEPQLSAALEYVRSHPLDPIDTVDFERECGVGVIVTPEQIEEAVEAAINRHRPQLLVERYHFNMGLLMGEARAVLKWADGKMIKNEV | P47897 | QARS_HPRR2960729 | PrEST |
| arginyl-tRNA synthetase | ArgRS | RARS1 | | | 1-660 | MDVLVSECSARLLQQEEEIKSLTAEIDRLKNCGCLGASPNLEQLQEENLKLKYRLNILRKSLQAERNKPTKNMINIISRLQEVFGHAIKAAYPDLENPPLLVTPSQQAKFGDYQCNSAMGISQMLKTKEQKVNPREIAENITKHLPDNECIEKVEIAGPGFINVHLRKDFVSEQLTSLLVNGVQLPALGENKKVIVDFSSPNIAKEMHVGHLRSTIIGESISRLFEFAGYDVLRLNHVGDWGTQFGMLIAHLQDKFPDYLTVSPPIGDLQVFYKESKKRFDTEEEFKKRAYQCVVLLQGKNPDITKAWKLICDVSRQELNKIYDALDVSLIERGESFYQDRMNDIVKEFEDRGFVQVDDGRKIVFVPGCSIPLTIVKSDGGYTYDTSDLAAIKQRLFEEKADMIIYVVDNGQSVHFQTIFAAAQMIGWYDPKVTRVFHAGFGVVLGEDKKKFKTRSGETVRLMDLLGEGLKRSMDKLKEKERDKVLTAEELNAAQTSVAYGCIKYADLSHNRLNDYIFSFDKMLDDRGNTAAYLLYAFTRIRSIARLANIDEEMLQKAARETKILLDHEKEWKLGRCILRFPEILQKILDDLFLHTLCDYIYELATAFTEFYDSCYCVEKDRQTGKILKVNMWRMLLCEAVAAVMAKGFDILGIKPVQRM | P54136 | RARS_M1-M660 | biotinylated recombinant protein |
| arginyl-tRNA synthetase | ArgRS | RARS1 | | | 443-584 | VVLGEDKKKFKTRSGETVRLMDLLGEGLKRSMDKLKEKERDKVLTAEELNAAQTSVAYGCIKYADLSHNRLNDYIFSFDKMLDDRGNTAAYLLYAFTRIRSIARLANIDEEMLQKAARETKILLDHEKEWKLGRCILRFPEI | P54136 | RARS_HPRR620009 | PrEST* |
| arginyl-tRNA synthetase | ArgRS | RARS1 | | | 215-364 | TIIGESISRLFEFAGYDVLRLNHVGDWGTQFGMLIAHLQDKFPDYLTVSPPIGDLQVFYKESKKRFDTEEEFKKRAYQCVVLLQGKNPDITKAWKLICDVSRQELNKIYDALDVSLIERGESFYQDRMNDIVKEFEDRGFVQVDDGRKIV | P54136 | RARS_HPRR620010 | PrEST |
| arginyl-tRNA synthetase | ArgRS | RARS1 | | | 70-660 | TKNMINIISRLQEVFGHAIKAAYPDLENPPLLVTPSQQAKFGDYQCNSAMGISQMLKTKEQKVNPREIAENITKHLPDNECIEKVEIAGPGFINVHLRKDFVSEQLTSLLVNGVQLPALGENKKVIVDFSSPNIAKEMHVGHLRSTIIGESISRLFEFAGYDVLRLNHVGDWGTQFGMLIAHLQDKFPDYLTVSPPIGDLQVFYKESKKRFDTEEEFKKRAYQCVVLLQGKNPDITKAWKLICDVSRQELNKIYDALDVSLIERGESFYQDRMNDIVKEFEDRGFVQVDDGRKIVFVPGCSIPLTIVKSDGGYTYDTSDLAAIKQRLFEEKADMIIYVVDNGQSVHFQTIFAAAQMIGWYDPKVTRVFHAGFGVVLGEDKKKFKTRSGETVRLMDLLGEGLKRSMDKLKEKERDKVLTAEELNAAQTSVAYGCIKYADLSHNRLNDYIFSFDKMLDDRGNTAAYLLYAFTRIRSIARLANIDEEMLQKAARETKILLDHEKEWKLGRCILRFPEILQKILDDLFLHTLCDYIYELATAFTEFYDSCYCVEKDRQTGKILKVNMWRMLLCEAVAAVMAKGFDILGIKPVQRM | P54136 | RARS_T70-M660 | biotinylated recombinant protein |
| seryl-tRNA synthetase | SerRS | SARS1 | | | 83-212 | NVLSFDDLTADALANLKVSQIKKVRLLIDEAILKCDAERIKLEAERFENLREIGNLLHPSVPISNDEDVDNKVERIWGDCTVRKKYSHVDLVVMVDGFEGEKGAVVAGSRGYFLKGVLVFLEQALIQYAL | P49591 | SARS_HPRR1390005 | PrEST |
| seryl-tRNA synthetase | SerRS | SARS1 | | | 216-303 | GSRGYIPIYTPFFMRKEVMQEVAQLSQFDEELYKVIGKGSEKSDDNSYDEKYLIATSEQPIAALHRDEWLRPEDLPIKYAGLSTCFRQ | P49591 | SARS_HPRR4160317 | PrEST |
| seryl-tRNA synthetase | SerRS | SARS1 | | | 1-514 | MVLDLDLFRVDKGGDPALIRETQEKRFKDPGLVDQLVKADSEWRRCRFRADNLNKLKNLCSKTIGEKMKKKEPVGDDESVPENVLSFDDLTADALANLKVSQIKKVRLLIDEAILKCDAERIKLEAERFENLREIGNLLHPSVPISNDEDVDNKVERIWGDCTVRKKYSHVDLVVMVDGFEGEKGAVVAGSRGYFLKGVLVFLEQALIQYALRTLGSRGYIPIYTPFFMRKEVMQEVAQLSQFDEELYKVIGKGSEKSDDNSYDEKYLIATSEQPIAALHRDEWLRPEDLPIKYAGLSTCFRQEVGSHGRDTRGIFRVHQFEKIEQFVYSSPHDNKSWEMFEEMITTAEEFYQSLGIPYHIVNIVSGSLNHAASKKLDLEAWFPGSGAFRELVSCSNCTDYQARRLRIRYGQTKKMMDKVEFVHMLNATMCATTRTICAILENYQTEKGITVPEKLKEFMPPGLQELIPFVKPAPIEQEPSKKQKKQHEGSKKKAAARDVTLENRLQNMEVTDA | P49591 | SARS_M1-A514 | biotinylated recombinant protein |
| U1 small nuclear ribonucleoprotein 70 kDa | | SNRNP 70 | U1RNP | MAA | 60-216 | AETREERMERKRREKIERRQQEVETELKMWDPHNDPNAQGDAFKTLFVARVNYDTTESKLRREFEVYGPIKRIHMVYSKRSGKPRGYAFIEYEHERDMHSAYKHADGKKIDGRRVLVDVERGRTVKGWRPRRLGGGLGGTRRGGADVNIRHSGRDDT | P08621 | SNRNP70_A60-T216 | biotinylated recombinant protein |
| U1 small nuclear ribonucleoprotein 70 kDa | | SNRNP 70 | U1RNP | MAA | 99-187 | GDAFKTLFVARVNYDTTESKLRREFEVYGPIKRIHMVYSKRSGKPRGYAFIEYEHERDMHSAYKHADGKKIDGRRVLVDVERGRTVKGW | P08621 | SNRNP70_HPRR3210218 | PrEST |

| | | | | | | | | | |
|---|---|---|---|---|---|---|---|---|---|
| U1 small nuclear ribonucleoprotein 70 kDa | | SNRNP 70 | U1RNP | MAA | 16-98 | RDPIPYLPPLEKLPHEKHHNQPYCGIAPYIREFEDPRDAPPPTRAETREERMERKRREKIERRQQEVETELKMWDPHNDPNAQ | P08621 | SNRNP70_HPRR3210219 | PrEST |
| Sjogren syndrome antigen B | | SSB | SSB | MAA | 1-408 | MAENGDNEKMAALEAKICHQIEYYFGDFNLPRDKFLKEQIKLDEGWVPLEIMIKFNRLNRLTTDFNVIVEALSKSKAELMEISEDKTKIRRSPSKPLPEVTDEYKNDVKNRSVYIKGFPTDATLDDIKEWLEDKGQVLNIQMRRTLHKAFKGSIFVVFDSIESAKKFVETPGQKYKETDLLILFKDDYFAKKNEERKQNKVEAKLRAKQEQEAKQKLEEDAEMKSLEEKIGCLLKFSGDLDDQTCREDLHILFSNHGEIKWIDFVRGAKEGIILFKEKAKEALGKAKDANNGNLQLRNKEVTWEVLEGEVEKEALKKIIEDQQESLNKWKSKGRRFKGKGKGNKAAQPGSGKGKVQFQGKKTKFASDDEHDEHDENGATGPVKRAREETDKEEPASKQQKTENGAGDQ | P05455 | SSB_M1-Q408 | biotinylated recombinant protein |
| Sjogren syndrome antigen B | | SSB | SSB | MAA | 15-126 | AKICHQIEYYFGDFNLPRDKFLKEQIKLDEGWVPLEIMIKFNRLNRLTTDFNVIVEALSKSKAELMEISEDKTKIRRSPSKPLPEVTDEYKNDVKNRSVYIKGFPTDATLDD | P05455 | SSB_HPRR2010045 | PrEST |
| Sjogren syndrome antigen B | | SSB | SSB | MAA | 156-217 | VVFDSIESAKKFVETPGQKYKETDLLILFKDDYFAKKNEERKQNKVEAKLRAKQEQEAKQKL | P05455 | SSB_HPRR2010046 | PrEST |
| threonyl-tRNA synthetase | ThrRS | TARS1 | PL7 | MSA | 1-723 | MFEEKASSPSGKMGGEEKPIGAGEEKQKEGGKKKNKEGSGDGGRAELNPWPEYIYTRLEMYNILKAEHDSILAEKAEKDSKPIKVTLPDGKQVDAESWKTTPYQIACGISQGLADNTVIAKVNNVVWDLDRPLEEDCTLELLKFEDEEAQAVYWHSSAHIMGEAMERVYGGCLCYGPPIENGFYYDMYLEEGGVSSNDFSSLEALCKKIIKEKQAFERLEVKKETLLAMFKYNKFKCRILNEKVNTPTTTVYRCGPLIDLCRGPHVRHTGKIKALKIHKNSSTYWEGKADMETLQRIYGISFPDPKMLKEWEKFQEEAKNRDHRKIGRDQELYFFHELSPGSCFFLPKGAYIYNALIEFIRSEYRKRGFQEVVTPNIFNSRLWMTSGHWQHYSENMFSFEVEKELFALKPMNCPGHCLMFDHRPRSWRELPLRLADFGVLHRNELSGALTGLTRVRRFQQDDAHIFCAMEQIEDEIKGCLDFLRTVYSVFGFSFKLNLSTRPEKFLGDIEVWDQAEKQLENSLNEFGEKWELNSGDGAFYGPKIDIQIKDAIGRYHQCATIQLDFQLPIRFNLTYVSHDGDDKKRPVIVHRAILGSVERMIAILTENYGGKWPFWLSPRQVMVVPVGPTCDEYAQKVRQQFHDAKFMADIDLDPGCTLNKKIRNAQLAQYNFILVVGEKEKISGTVNIRTRDNKVHGERTISETIERLQQLKEFRSKQAEEEF | P26639 | TARS_M1-F723 | biotinylated recombinant protein |
| threonyl-tRNA synthetase | ThrRS | TARS1 | PL7 | MSA | 1-64 | MFEEKASSPSGKMGGEEKPIGAGEEKQKEGGKKKNKEGSGDGGRAELNPWPEYIYTRLEMYNIL | P26639 | TARS_HPRR3010092 | PrEST |
| threonyl-tRNA synthetase | ThrRS | TARS1 | PL7 | MSA | 691-723 | RDNKVHGERTISETIERLQQLKEFRSKQAEEEF | P26639 | TARS_HPRR3890005 | PrEST |
| threonyl-tRNA synthetase | ThrRS | TARS1 | PL7 | MSA | 320-723 | NRDHRKIGRDQELYFFHELSPGSCFFLPKGAYIYNALIEFIRSEYRKRGFQEVVTPNIFNSRLWMTSGHWQHYSENMFSFEVEKELFALKPMNCPGHCLMFDHRPRSWRELPLRLADFGVLHRNELSGALTGLTRVRRFQQDDAHIFCAMEQIEDEIKGCLDFLRTVYSVFGFSFKLNLSTRPEKFLGDIEVWDQAEKQLENSLNEFGEKWELNSGDGAFYGPKIDIQIKDAIGRYHQCATIQLDFQLPIRFNLTYVSHDGDDKKRPVIVHRAILGSVERMIAILTENYGGKWPFWLSPRQVMVVPVGPTCDEYAQKVRQQFHDAKFMADIDLDPGCTLNKKIRNAQLAQYNFILVVGEKEKISGTVNIRTRDNKVHGERTISETIERLQQLKEFRSKQAEEEF | P26639 | TARS_N320-F723 | biotinylated recombinant protein |
| tripartite motif containing 21 | | TRIM21 | SSA (Ro-52) | MAA | 200-239 | LEKDEREQLRILGEKEAKLAQQSQALQELISELDRRCHSS | P19474 | TRIM21_HPRR1060031 | PrEST |
| E3 ubiquitin-protein ligase TRIM33 | | TRIM33 | TIF1-y | MSA | 882-1127 | DDDPNEDWCAVCQNGGDLLCCEKCPKVFHLTCHVPTLLSFPSGDWICTFCRDIGKPEVEYDCDNLQHSKKGKTAQGLSPVDQRKCERLLLYLYCHELSIEFQEPVPASIPNYYKIIKKPMDLSTVKKKLQKKHSQHYQIPDDFVADVRLIFKNCERFNEMMKVVQVYADTQEINLKADSEVAQAGKAVALYFEDKLTEIYSDRTFAPLPEFEQEEDDGEVTEDSDEDFIQPRRKRLKSDERPVHIK | Q9UPN9 | TRIM33_D882-K1127 | biotinylated recombinant protein |
| TROVE domain family member 2 | | TROVE 2 | SSA (Ro-60) | MAA | 3-145 | ESVNQMQPLNEKQIANSQDGYVWQVTDMNRLHRFLCFGSEGGTYYIKEQKLGLENAEALIRLIEDGRGCEVIQEIKSFSQEGRTTKQEPMLFALAICSQCSDISTKQAAFKAVSEVCRIPTHLFTFIQFKKDLKESMKCGMWG | P10155 | TROVE2_HPRR400060 | PrEST* |
| TROVE domain family member 2 | | TROVE 2 | SSA (Ro-60) | MAA | 371-506 | FLLAVDVSASMNQRVLGSILNASTVAAAMCMVVTRTEKDSYVVAFSDEMVPCPVTTDMTLQQVLMAMSQIPAGGTDCSLPMIWAQKTNTPADVFIVFTDNETFAGGVHPAIALREYRKKMDIPAKLIVCGMTSNGF | P10155 | TROVE2_HPRR400061 | PrEST |
| valyl-tRNA synthetase | ValRS | VARS1 | | | 1074-1169 | RLPRRMPQAPPSLCVTPYPEPSECSWKDPEAEAALELALSITRAVRSLRADYNLTRIRPDCFLEVADEATGALASAVSGYVQALASAGVVAVLALG | P26640 | VARS_HPRR3340369 | PrEST |
| valyl-tRNA synthetase | ValRS | VARS1 | | | 1171-1264 | AVALASDRCSIHLQLQGLVDPARELGKLQAKRVEAQRQAQRLRERRAASGYPVKVPLEVQEADEAKLQQTEAELRKVDEAIALFQKML | P26640 | VARS_HPRR3340371 | PrEST |
| valyl-tRNA synthetase | ValRS | VARS1 | | | 1-300 | MSTLYVSPHPDAFPSLRALIAARYGEAGEGPGWGGAHPRICLQPPPTSRTSFPPPRLPALEQGPGGLWVWGATAVAQLLWPAGLGGPGGSRAAVLVQQWVSYADTELIPAACGATLPALGLRSSAQDPQAVLGALGRALSPLEEWLRLHTYLAGEAPTLADLAAVTALLLPFRYVLDPPARRIWNNVTRWFVTCVRQPEFRAVLGEVVLYSGARPLSHQPGPEAPALPKTAAQLKKEAKKREKLEKFQQKQKIQQQQPPPGEKKPKPEKREKRDPGVITYDLPTPPGEKKDVSGPMPDSY | P26640 | VARS_M1-Y300 | biotinylated recombinant protein* |
| tryptophanyl-tRNA synthetase | TrpRS | WARS1 | | | 270-405 | TDSDCIGKISFPAIQAAPSFSNSFPQIFRDRTDIQCLIPCAIDQDPYFRMTRDVAPRIGYPKPALLHSTFFPALQGAQTKMSASDPNSSIFLTDTAKQIKTKVNKHAFSGGRDTIEEHRQFGGNCDVDVSFMYLTF | P23381 | WARS_HPRR141609 | PrEST* |
| tryptophanyl-tRNA synthetase | TrpRS | WARS1 | | | 76-219 | DATEAEEDFVDPWTVQTSSAKGIDYDKLIVRFGSSKIDKELINRIERATGQRPHHFLRRGIFFSHRDMNQVLDAYENKKPFYLYTGRGPSSEAMHVGHLIPFIFTKWLQDVFNVPLVIQMTDDEKYLWKDLTLDQAYSYAVENA | P23381 | WARS_HPRR141610 | PrEST |
| tryptophanyl-tRNA synthetase | TrpRS | WARS1 | | | 381-405 | RDTIEEHRHFGGNCVVDVSFMYLTF | P23381 | WARS_HPRR3660104 | PrEST |
| tryptophanyl-tRNA synthetase | TrpRS | WARS1 | | | 1-471 | MPNSEPASLLELFNSIATQGELVRSLKAGNASKDEIDSAVKMLVSLKMSYKAAAGEDYKADCPPGNPAPTSNHGPDATEAEEDFVDPWTVQTSSAKGIDYDKLIVRFGSSKIDKELINRIERATGQRPHHFLRRGIFFSHRDMNQVLDAYENKKPFYLYTGRGPSSEAMHVGHLIPFIFTKWLQDVFNVPLVIQMTDDEKYLWKDLTLDQAYSYAVENAKDIIACGFDINKTFIFSDLDYMGMSSGFYKNVVKIQKHVTFNQVKGIFGFTDSDCIGKISFPAIQAAPSFSNSFPQIFRDRTDIQCLIPCAIDQDPYFRMTRDVAPRIGYPKPALLHSTFFPALQGAQTKMSASDPNSSIFLTDTAKQIKTKVNKHAFSGGRDTIEEHRQFGGNCDVDVSFMYLTFFLEDDDKLEQIRKDYTSGAMLTGELKKALIEVLQPLIAEHQARRKEVTDEIVKEFMTPRKLSFDFQ | P23381 | WARS_M1-Q471 | biotinylated recombinant protein |
| X-ray repair cross complementing 6 | | XRCC6 | Ku | MAA | 407-506 | PPYFVALVPQEEELDDQKIQVTPPGFQLVFLPFADDKRKMPFTEKIMATPEQVGKMKAIVEKLRFTYRSDSFENPVLQQHFRNLEALALDLMEPEQAVDL | P12956 | XRCC6_HPRR3530014 | PrEST |

| | | | | | | | | | |
|---|---|---|---|---|---|---|---|---|---|
| X-ray repair cross complementing 6 | | XRCC6 | Ku | MAA | 156-252 | DVQFKMSHKRIMLFTNEDNPHGNDSAKASRARTKAGDLRDTGIFLDLMHLKKPGGFDISLFYRDIISIAEDEDLRVHF EESSKLEDLLRKVRAKETR | P12956 | XRCC6_HPRR3530017 | PrEST |
| tyrosyl-tRNA synthetase | TyrRS | YARS1 | HA | MSA | 1-528 | MGDAPSPEEKLHLITRNLQEVLGEEKLKEILKERELKIYWGTATTGKPHVAYFVPMSKIADFLKAGCEVTILFADLH AYLDNMKAPWELLELRVSYYENVIKAMLESIGVPLEKLKFIKGTDYQLSKEYTLDVYRLSSVVTQHDSKKAGAEVV KQVEHPLLSGLLYPGLQALDEEYLKVDAQFGGIDQRKIFTFAEKYLPALGYSKRVHLMNPMVPGLTGSKMSSSEEES KIDLLDRKEDVKKKLKKAFCEPGNVENNGVLSFIKHVLFPLKSEFVILRDEKWGGNKTYTAYVDLEKDFAAEVVHP GDLKNSVEVALNKLLDPIREKFNTPALKKLASAAYPDPSKQKPMAKGPAKNSEPEEVIPSRLDIRVGKIITVEKHPDA DSLYVEKIDVGEAEPRTVVSGLVQFVPKEELQDRLVVVLCNLKPQKMRGVESQGMLLCASIEGINRQVEPLDPPAGS APGEHVFVKGYEKGQPDEELKPKKKVFEKLQADFKISEECIAQWKQTNFMTKLGSISCKSLKGGNIS | P54577 | YARS_M1-S528 | biotinylated recombinant protein |
| tyrosyl-tRNA synthetase | TyrRS | YARS1 | HA | MSA | 1-341 | MGDAPSPEEKLHLITRNLQEVLGEEKLKEILKERELKIYWGTATTGKPHVAYFVPMSKIADFLKAGCEVTILFADLH AYLDNMKAPWELLELRVSYYENVIKAMLESIGVPLEKLKFIKGTDYQLSKEYTLDVYRLSSVVTQHDSKKAGAEVV KQVEHPLLSGLLYPGLQALDEEYLKVDAQFGGIDQRKIFTFAEKYLPALGYSKRVHLMNPMVPGLTGSKMSSSEEES KIDLLDRKEDVKKKLKKAFCEPGNVENNGVLSFIKHVLFPLKSEFVILRDEKWGGNKTYTAYVDLEKDFAAEVVHP GDLKNSVEVALNKLLDPIREKFNTPALKKLASAAY | P54577 | YARS_M1-Y341 | biotinylated recombinant protein |
| tyrosyl-tRNA synthetase | TyrRS | YARS1 | HA | MSA | 126-243 | SKEYTLDVYRLSSVVTQHDSKKAGAEVVKQVEHPLLSGLLYPGLQALDEEYLKVDAQFGGIDQRKIFTFAEKYLPA LGYSKRVHLMNPMVPGLTGSKMSSSEEESKIDLLDRKEDVKK | P54577 | YARS_HPRR2320094 | PrEST |
| tyrosyl-tRNA synthetase | TyrRS | YARS1 | HA | MSA | 253-385 | GNVENNGVLSFIKHVLFPLKSEFVILRDEKWGGNKTYTAYVDLEKDFAAEVVHPGDLKNSVEVALNKLLDPIREKF NTPALKKLASAAYPDPSKQKPMAKGPAKNSEPEEVIPSRLDIRVGKIITVEKHPDAD | P54577 | YARS_HPRR2320095 | PrEST |
| tyrosyl-tRNA synthetase | TyrRS | YARS1 | HA | MSA | 422-518 | VLCNLKPQKMRGVESQGMLLCASIEGINRQVEPLDPPAGSAPGEHVFVKGYEKGQPDEELKPKKKVFEKLQADFKIS EECIAQWKQTNFMTKLGSIS | P54577 | YARS_HPRR2320096 | PrEST* |
| tyrosyl-tRNA synthetase | TyrRS | YARS1 | HA | MSA | 5-119 | PSPEEKLHLITRNLQEVLGEEKLKEILKERELKIYWGTATTGKPHVAYFVPMSKIADFLKAGCEVTILFADLHAYLDN MKAPWELLELRVSYYENVIKAMLESIGVPLEKLKFIK | P54577 | YARS_HPRR2320097 | PrEST* |

*All above stated antigens were included in the multiplex bead array analysis of plasma. The antigens marked with * were removed in the analysis of serum, and the remaining antigens were included. MAA, myositis-associated autoantibody; MSA, myositis-specific autoantibody; PrEST, protein epitope signature tag.*
