## Supplementary Information for "Autoantigenic properties of the aminoacyl tRNA synthetase family in idiopathic inflammatory myopathies"

#### SUPPLEMENTARY METHODS

##### Patients and population controls

Information on laboratory and clinical data was retrieved by revising the medical charts, the Swedish Rheumatology Quality Register for IIM (Swemyonet) [1] and the Euromyositis register [2]. Muscle involvement was based on the presence of at least one of the following features: myopathic weakness (manual muscle test-8 (MMT-8) below 80 and/or impaired muscle endurance by myositis functional index-2 [3]), muscle enzymes elevation (creatine kinase (CK), lactate dehydrogenase (LD), aspartate aminotransferase (ASAT), alanine aminotransferase (ALAT)), myopathic electromyography (EMG), pathological muscle biopsy consistent with myositis. One of the following skin manifestations had to be present to define the skin involvement: periungual erythema, mechanic's hand, Gottron's sign, Gottron's papules, V-sign, shawl sign, erythroderma, periorbital edema, heliotrope rash, calcinosis. Myositis specific autoantibodies (MSAs (anti Jo-1, -PL7, -PL12, -OJ, -EJ, -Mi-2, -SRP, -TIF1γ and -MDA5)) and/or myositis associated autoantibodies (MAAs (anti-U1RNP, -Pm-Scl, -Ku, -SSA, -SSB)) were analyzed using any of the following assays: immunoprecipitation, Line Blot (Euroimmun), or ELISA). With regards to anti-SSA antibodies, information on reactivity to Ro52 (TRIM21) and Ro60 (TROVE2) was not available for all patients and therefore was not reported. Autoantibody positivity and clinical manifestations (myositis, skin pathology, arthritis, Raynaud's phenomenon, dysphagia, ILD, cardiac involvement (any of myocarditis, pericarditis or arrythmia), cancer diagnosis was assigned to patients if ever confirmed during the follow-up (interval time occurring between the time of diagnosis and the time of the last visit at the Rheumatology Clinic). Other autoimmune diseases included: rheumatoid arthritis, systemic lupus erythematosus, systemic sclerosis, morphea, sjögren syndrome, mixed connective tissue disease.

##### ELISA

Streptavidin-coated (2 µg/ml diluted in PBS) 384 well plates were incubated over night at 4°C. The next day, the plates were washed four times in PBS-T (0.05% (v/v) Tween-20) and blocked for 2 h at room temperature with 50 µl blocking buffer (PBS with 0.05% (v/v) Tween-20 and 5 % (w/v) BSA). Biotinylated recombinant protein was added at 1 µg/ml diluted in PBT (PBS with 0.05% (v/v) Tween-20 and 0.5 % (w/v) BSA), 25 µl/well before incubation for 1 h. Plates were washed four times with PBS-T and plasma diluted (in a five-fold dilution series from 25x to 16500x dilution) and incubated for 1 h in dilution buffer (PBS with 0.05% (v/v) Tween-20 and 1 % (w/v) BSA) were added in duplicates (25 µl/well) and let to incubate for 1.5 h. Subsequently, plates were washed four times in PBS-T and incubated with horseradish peroxidase conjugated rabbit anti-human IgG antibody (Dako #P0214), diluted 1:4000, in PBT for 1h at RT. After washing four times, 25 µl/well TMB substrate (Sigma Aldrich) was added to the plates and the reaction was stopped after 3 min by adding 25 µl/well of 1 M $H_2SO_4$. The optical density (OD) was read at 450 nm with 620 nm background removal using a microplate reader (SpectraMaxPlus384) and SoftMax Pro 6.5 software (Molecular Devices). All samples were tested against MDA5 as a reference antigen, and streptavidin (SA) as background. In addition, sample from a patient with anti-MDA5 autoantibodies was used as control plasma for background signals of the specific autoantigen (Supplementary Fig. 11).

### SUPPLEMENTARY RESULTS

**Supplementary Table 1:** Demographics of patient and population control samples

| | IIM n=217 | PC (n=156) |
|---|---|---|
| Age at sample, mean years (SD) | 59.2 (14.8) | 53.9 (15.4) |
| Sex, n (%) women | 137 (63.1) | 103 (64.8) |

IIM, Idiopathic inflammatory myopathies; PC, population control.

**Supplementary Table 2:** Number of patients with antibodies (IgG) targeting any of the versions of the specific antigen tested for, for each cut-off analyzed.

| Antigen | MADs>50<br>n (%) | MADs>100<br>n (%) | MADs>150<br>n (%) | MADs>200<br>n (%) |
|---|---|---|---|---|
| HisRS (Jo1) | 47 (21.7) | 43 (19.8) | 41 (18.9) | 41 (18.9) |
| ThrRS (PL7) | 13 (6.0) | 9 (4.1) | 6 (2.8) | 6 (2.8) |
| AlaRS (PL12) | 6 (2.8) | 3 (1.4) | 2 (0.9) | 1 (0.5) |
| GlyRS (EJ) | 7 (3.3) | 2 (0.9) | 1 (0.5) | 1 (0.5) |
| IleRS (OJ) | 1 (0.5) | 0 (0.0) | 0 (0.0) | 0 (0.0) |
| AsnRS (KS) | 4 (1.8) | 2 (0.9) | 0 (0.0) | 0 (0.0) |
| PheRS (Zo) | 11 (5.1) | 5 (2.3) | 3 (1.4) | 2 (0.9) |
| TyrRS (HA) | 7 (3.2) | 3 (1.4) | 2 (0.9) | 2 (0.9) |
| LysRS (Sc) | 2 (0.9) | 1 (0.5) | 1 (0.5) | 0 (0.0) |
| GlnRS (JS) | 4 (1.8) | 2 (0.9) | 0 (0.0) | 0 (0.0) |
| TrpRS (WARS) | 1 (0.5) | 1 (0.5) | 0 (0.0) | 0 (0.0) |
| SerRS | 1 (0.5) | 1 (0.5) | 1 (0.5) | 1 (0.5) |
| EPRS | 2 (0.9) | 1 (0.5) | 1 (0.5) | 1 (0.5) |
| ArgRS | 6 (2.8) | 3 (1.4) | 2 (0.9) | 1 (0.5) |
| MetRS | 1 (0.5) | 1 (0.5) | 0 (0.0) | 0 (0.0) |
| LeuRS | 1 (0.5) | 0 (0.0) | 0 (0.0) | 0 (0.0) |
| ValRS | 8 (3.7) | 2 (0.9) | 1 (0.5) | 1 (0.5) |
| CysRS | 3 (1.4) | 2 (0.9) | 1 (0.5) | 1 (0.5) |
| AspRS | 1 (0.5) | 0 (0.0) | 0 (0.0) | 0 (0.0) |
| AIMP1 | 13 (6.0) | 8 (3.7) | 6 (2.8) | 5 (2.3) |
| AIMP2 | 5 (2.3) | 2 (0.9) | 2 (0.9) | 1 (0.5) |
| AIMP3 | 1 (0.5) | 0 (0.0) | 0 (0.0) | 0 (0.0) |
| Mi-2 | 17 (7.8) | 6 (2.8) | 3 (1.4) | 1 (0.5) |
| MDA5 | 11 (5.1) | 8 (3.7) | 7 (3.2) | 6 (2.8) |
| TRIM33 (TIF1γ) | 9 (4.1) | 6 (2.8) | 4 (1.8) | 3 (1.4) |
| SSA | 66 (30.4) | 56 (25.8) | 52 (24.0) | 44 (20.3) |
| SSB | 30 (13.8) | 24 (11.1) | 17 (7.8) | 13 (6.0) |
| U1 RNP | 16 (7.4) | 12 (5.5) | 8 (3.7) | 7 (3.2) |
| Ku | 0 (0.0) | 0 (0.0) | 0 (0.0) | 0 (0.0) |
| Pm-Scl | 16 (7.4) | 11 (5.1) | 8 (3.7) | 7 (3.2) |

The result from this analysis was included in the decision of selecting MADs>100 as an appropriate cut-off for this study. MADs, median absolute deviations; AIMP1-3, aaRS complex interacting multifunctional protein 1-3; EJ, GlyRS; HA, TyrRS; Jo1, HisRS; KS, AsnRS; Ku, X-ray repair cross complementing (XRCC) 6; MDA5, interferon-induced helicase C domain-containing protein 1; Mi-2, chromatin organization modifier helicase (CHD) 3 and 4; OJ, IleRS; PL7, ThrRS; PL12, AlaRS; Pm-Scl, polymyositis-scleroderma overlap syndrome-associated antigen 75 (exosome component 9) and 100 (exosome component 10); SSA, Ro52 (tripartite motif containing 21(TRIM21)) and Ro60 (TROVE domain family member 2 (TROVE2)); SSB, Sjogren syndrome antigen B; TIF1-γ, E3 ubiquitin-protein ligase (TRIM33); U1RNP, small nuclear ribonucleoprotein U1 subunit 70; Zo, PheRS.

**Supplementary Table 3:** Patients positive for more than one anti-aaRS or one anti-aaRS in combination with a known myositis specific autoantibody. The autoantigen for the specific antibody is stated in the table.

| Patient | Known antibody positivity | Reactivities detected in this study | Sequence similarity of double positive antigens (%) |
|---|---|---|---|
| 10 | HisRS (Jo1) | HisRS (Jo1), ThrRS (PL7) | 23.5 |
| 54 | HisRS (Jo1) | HisRS (Jo1), ThrRS (PL7) | 23.5 |
| 67 | | LysRS, GlnRS, ArgRS | 4.8, 16.9, 6.1 |
| 73 | HisRS (Jo1) | HisRS (Jo1), Mi-2α | 10.1 |
| 82 | | ThrRS (PL7), TIF1γ | 13.9 |
| 86 | HisRS (Jo1) | HisRS (Jo1), ArgRS | 21.4 |
| 96 | TIF1γ | TIF1γ, AlaRS (PL12) | 15.5 |
| 138 | | AlaRS (PL12), AsnRS (KS) | 6.7 |
| 150 | HisRS (Jo1) | HisRS (Jo1), Mi-2β | 8.6 |
| 155 | HisRS (Jo1) | HisRS (Jo1), AsnRS (KS) | 14.2 |
| 158 | TIF1γ | ThrRS (PL7), TIF1γ | 13.9 |
| 175 | AlaRS (PL12) | AlaRS (PL12), ThrRS (PL7) | 1.4 |
| 177 | HisRS (Jo1) | HisRS (Jo1), ArgRS | 16.1 |

The sequence similarities were calculated using pairwise sequence alignment (EMBOSS Needle). The protein versions that the patients were positive for were included in the alignment (tags and signal peptides excluded) and the highest similarity for that specific protein is reported. aaRS, aminoacyl tRNA synthetase; TIF1-γ, E3 ubiquitin-protein ligase (TRIM33); Mi-2, chromatin organization modifier helicase (CHD) 3 and 4.

**Supplementary Table 4:** Patients (n=12) positive for anti-Jo1, -PL7, -PL12, or -EJ, that were previously not known as positive for these autoantibodies. Nine patients are from the non-ASSD group. The autoantigen for the specific antibody is stated in the table.

| Patient | Group | Known antibody positivity | Reactivities detected in this study | Validated by ELISA |
|---|---|---|---|---|
| 5 | non-ASSD | seroneg | ThrRS (PL7) | |
| 10 | ASSD | HisRS (Jo1) | HisRS (Jo1), ThrRS (PL7) | |
| 53 | non-ASSD | seroneg | HisRS (Jo1) | |
| 54 | ASSD | HisRS (Jo1) | HisRS (Jo1), ThrRS (PL7) | |
| 82 | non-ASSD | seroneg | ThrRS (PL7) | |
| 96 | non-ASSD | TIF1γ | AlaRS (PL12) | |
| 122 | non-ASSD | seroneg | GlyRS (EJ) | |
| 138 | non-ASSD | seroneg | AlaRS (PL12), AsnRS (KS) | Supplementary Fig. 10 |
| 148 | non-ASSD | TIF1γ | HisRS (Jo1) | |
| 158 | non-ASSD | TIF1γ | ThrRS (PL7) | |
| 175 | ASSD | AlaRS (PL12) | AlaRS (PL12), ThrRS (PL7) | |
| 185 | non-ASSD | TIF1γ | ThrRS (PL7) | |

ASSD, anti-synthetase syndrome; seroneg, previously no myositis specific autoantibodies detected; TIF1γ, E3 ubiquitin-protein ligase TRIM33.

**Supplementary Table 5:** Patients positive for anti-AIMP1 and anti-AIMP2. The autoantigen for the specific antibody is stated in the table.

| Patient | Group | Known antibody positivity | Reactivities detected in this study | Validated by ELISA |
|---|---|---|---|---|
| 15 | ASSD | HisRS (Jo1) | AIMP2 | Supplementary Fig. 9 |
| 26 | non-ASSD | seroneg | TyrRS (HA), AIMP1 | Supplementary Fig. 9 |
| 70 | non-ASSD | seroneg | AIMP1 | |
| 95 | ASSD | IleRS (OJ) | AIMP1 | |
| 96 | non-ASSD | TIF1γ | AIMP2, TIF1γ, AlaRS (PL12) | |
| 110 | non-ASSD | seroneg | AIMP1 | |
| 124 | non-ASSD | MDA5 | MDA5, AIMP1 | |
| 138 | non-ASSD | seroneg | AlaRS (PL12), AsnRS (KS) AIMP1 | |
| 185 | non-ASSD | TIF1γ | ThrRS (PL7), AIMP1 | |
| 200 | ASSD | HisRS (Jo1) | HisRS (Jo1), AIMP1 | |

AIMP1 and 2, aaRS complex interacting multifunctional protein 1 and 2; ASSD, anti-synthetase syndrome; seroneg, previously no myositis specific autoantibodies detected. TIF1γ, E3 ubiquitin-protein ligase TRIM33; MDA5, interferon-induced helicase C domain-containing protein.

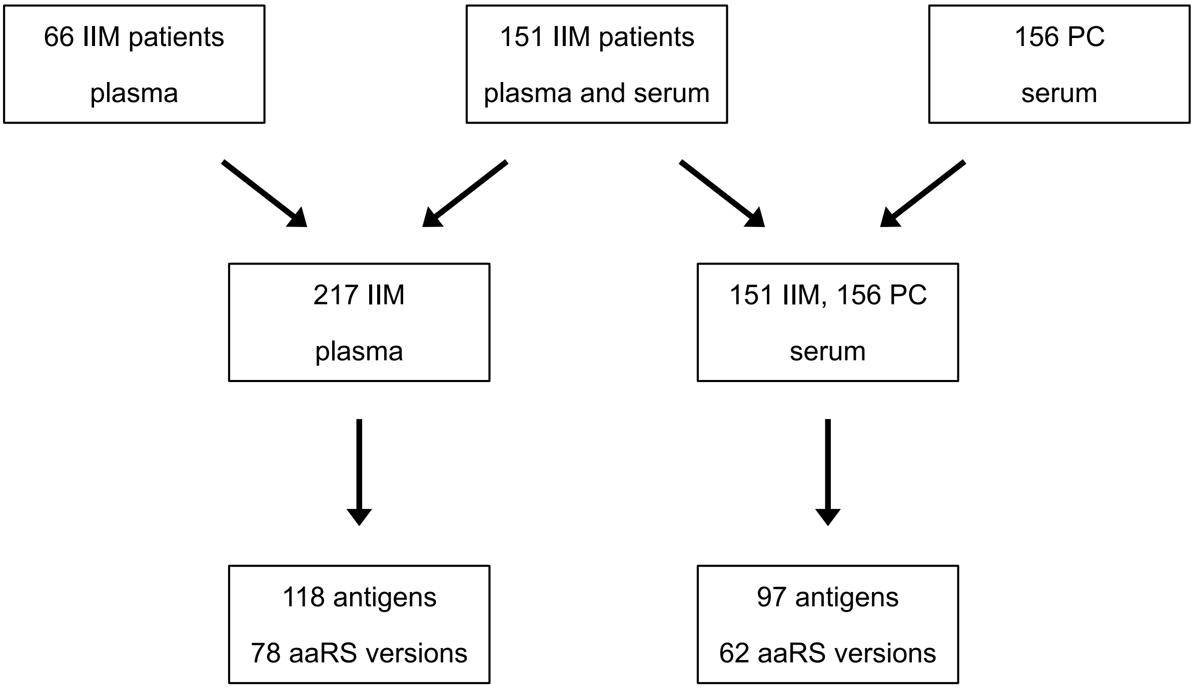


**Supplementary Fig. 1.** Experimental design of the study. 217 IIM plasma samples were analyzed using the multiplex bead array assay against 118 antigens (Supplementary Data). For 151 of the 217 IIM patients, serum was available and analyzed together with serum from 156 PC using the multiplex bead array assay against 97 antigens (Supplementary Data). IIM, idiopathic inflammatory myopathies, PC, population control; aaRS, aminoacyl tRNA synthetase.

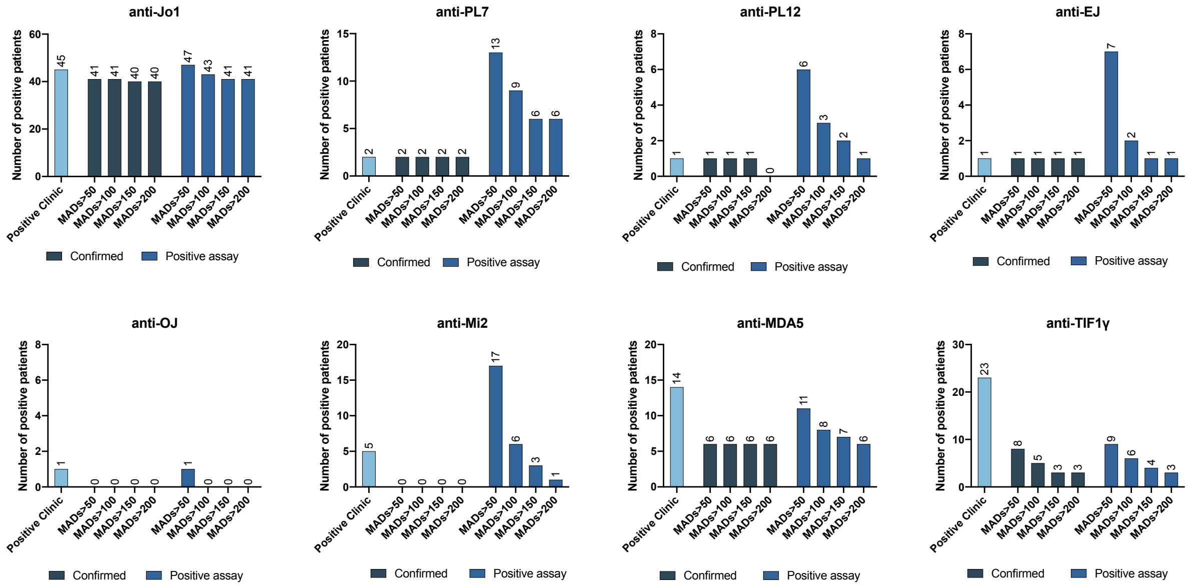


**Supplementary Fig. 2.** Patients positive for each of the myositis specific autoantibodies in previous tests (Positive Clinic, light blue), that could be confirmed in the multiplex bead array assay (Confirmed, dark blue), and total number of positive in the multiplex bead array assay (Positive assay, blue). EJ, GlyRS; Jo1, HisRS; MDA5, interferon-induced helicase C domain-containing protein 1; Mi-2, chromatin organization modifier helicase (CHD) 3 and 4; OJ, IleRS; PL7, ThrRS; PL12, AlaRS; TIF1-γ, E3 ubiquitin-protein ligase (TRIM33).

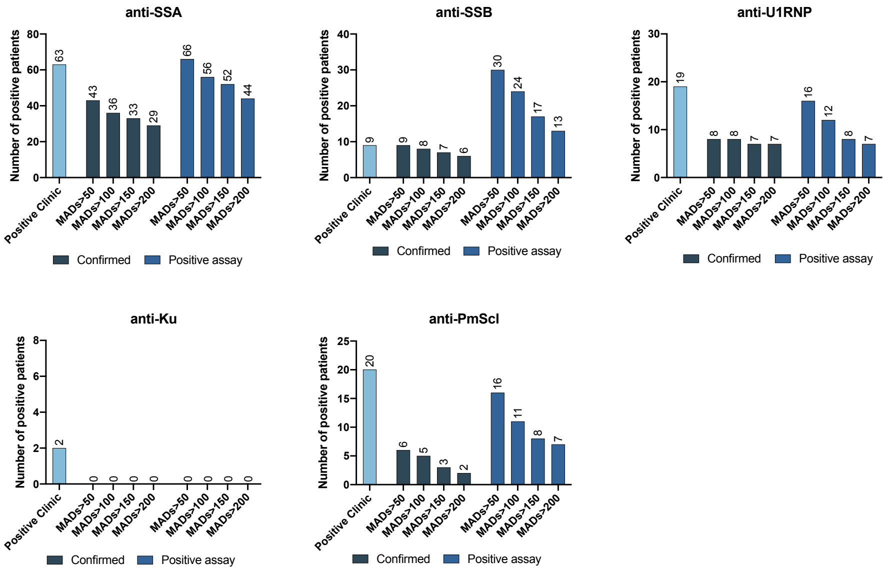


**Supplementary Fig. 3.** Patients positive for each of the myositis associated autoantibodies, in previous tests (Positive Clinic, light blue), that could be confirmed in the multiplex bead array assay (Confirmed, dark blue), and total number of positive in the multiplex bead array assay (Positive assay, blue). All 56 patients positive for anti-SSA in the assay were positive for anti-Ro52, none for anti-Ro60. SSA, Ro52 (tripartite motif containing 21(TRIM21)) and Ro60 (TROVE domain family member 2 (TROVE2)); SSB, Sjogren syndrome antigen B; U1RNP, small nuclear ribonucleoprotein U1 subunit 70; Ku, X-ray repair cross complementing (XRCC) 6; Pm-Scl, polymyositis-scleroderma overlap syndrome-associated antigen 75 (exosome component 9) and 100 (exosome component 10).

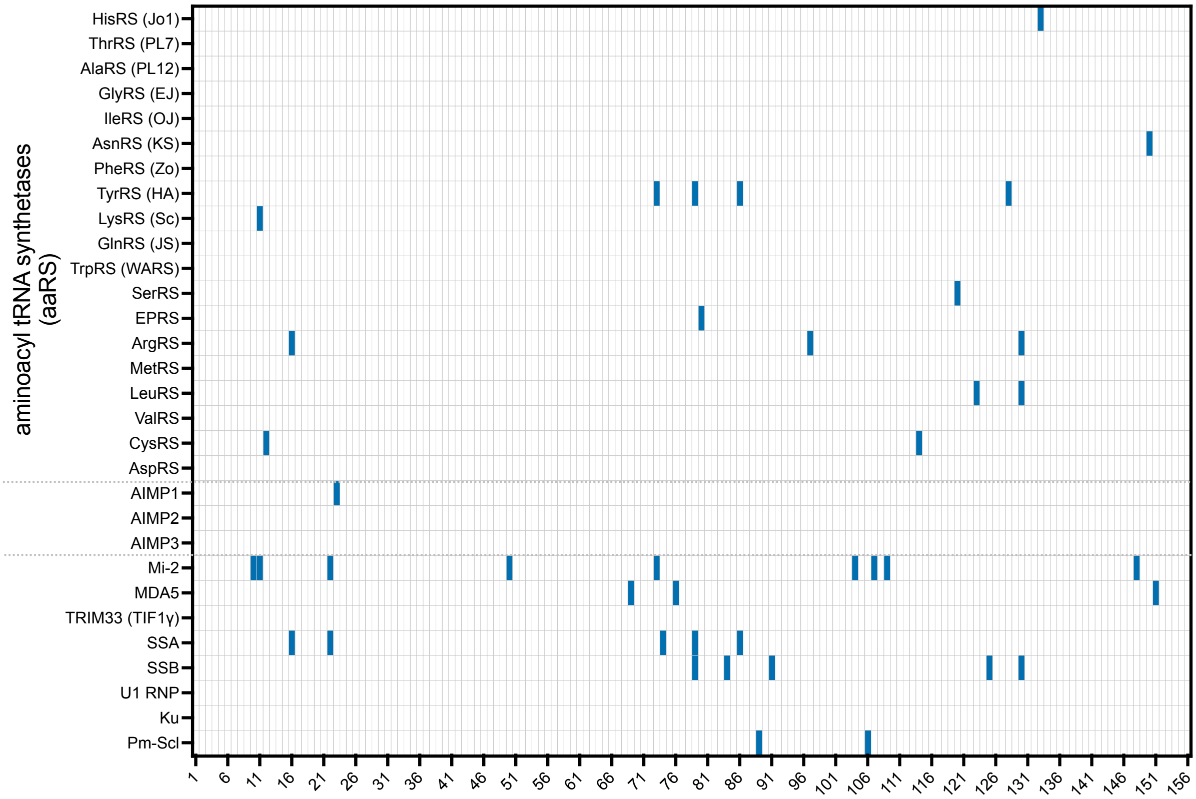


**Supplementary Fig. 4.** Autoantibody reactivities for the population control (PC) cohort against a panel of 30 antigens. Each column represents one individual, and each row represents one potential autoantigen. Reactivity was assigned positive (blue) if the criteria as defined in the method section were met for at least one of the included versions of a particular protein antigen. All cytoplasmic aaRS proteins are displayed above the dotted gray lines, the AIMP proteins are in between dotted lines and below are the additional myositis related proteins included in the study. AIMP1-3, aaRS complex interacting multifunctional protein 1-3; EJ, GlyRS; HA, TyrRS; Jo1, HisRS; KS, AsnRS; Ku, X-ray repair cross complementing (XRCC) 6; MDA5, interferon-induced helicase C domain-containing protein 1; Mi-2, chromatin organization modifier helicase (CHD) 3 and 4; OJ, IleRS; PL7, ThrRS; PL12, AlaRS; Pm-Scl, polymyositis-scleroderma overlap syndrome-associated antigen 75 (exosome component 9) and 100 (exosome component 10); SSA, Ro52 (tripartite motif containing 21 (TRIM21)) and Ro60 (TROVE domain family member 2 (TROVE2)); SSB, Sjogren syndrome antigen B; TIF1-γ, E3 ubiquitin-protein ligase (TRIM33); U1RNP, small nuclear ribonucleoprotein U1 subunit 70; Zo, PheRS.

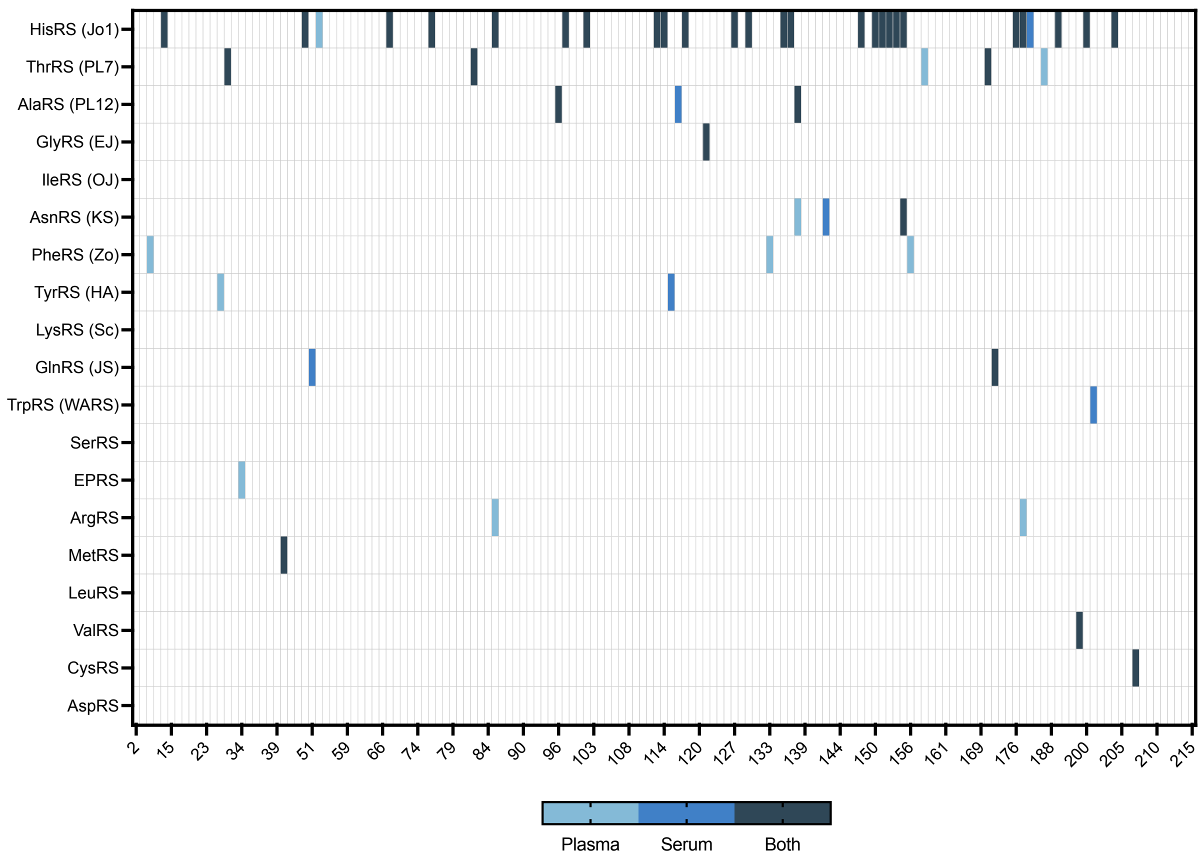


**Supplementary Fig. 5.** Autoantibody reactivities against the nineteen aaRS proteins for the 151 IIM patients with both serum and plasma availabe to control for sample differences. We found the same reactivities in both samples for 134/151 patients (89%). Reactivities could not be confirmed due to missing antigen in four patients while in eleven patients we observed an increased signal which did not reach the cut-off. In two patients (1.3%) reactivities did not correlate between plasma and serum samples (both targeting on of the ArgRS versions). Each column represents one individual, and each row represents one potential autoantigen. Reactivity was assigned positive if the criteria as defined in the method section were met for at least one of the included versions of a particular protein antigen. Light blue represents positive signal in plasma sample, blue in serum sample, and dark blue when displayed positive in both plasma and serum. EJ, GlyRS; HA, TyrRS; Jo1, HisRS; KS, AsnRS; OJ, IleRS; PL7, ThrRS; PL12, AlaRS; Zo, PheRS.

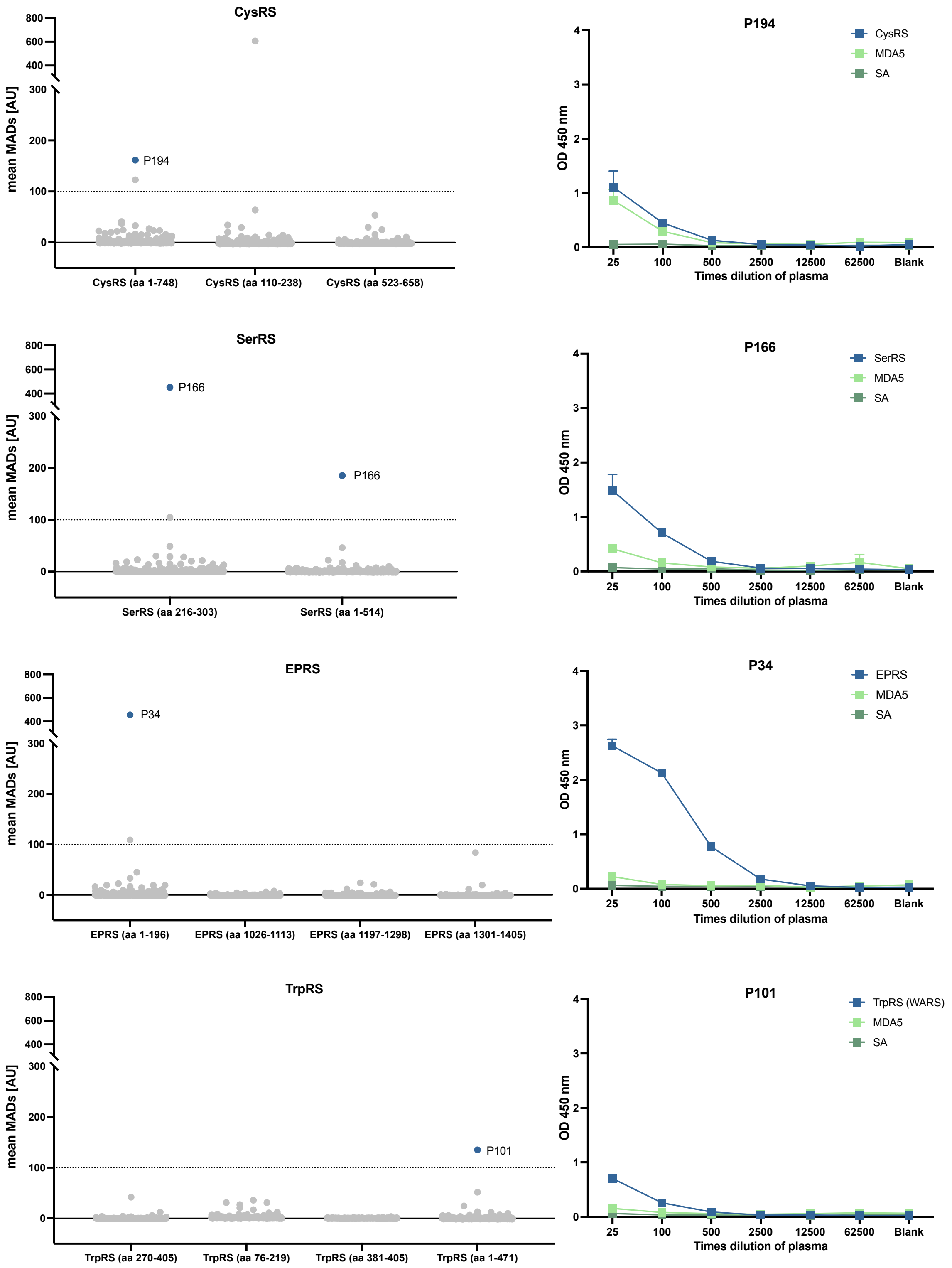


**Supplementary Fig. 6.** Validation of autoantibody reactivities. Results from the multiplex bead array assay (left) and ELISA (right) for patients with IgG targeting CysRS, SerRS, EPRS and TrpRS. The amino acid coverage for each antigen used in the multiplex bead array assay is displayed on the x-axis (left). The antigens used in ELISA were CysRS (aa 1-748), EPRS (aa 1-196), and TrpRS (aa 1-471). aa, amino acid; SA, streptavidin. MDA5, interferon-induced helicase C domain-containing protein 1; MADs, median absolute deviations; OD, optical density.

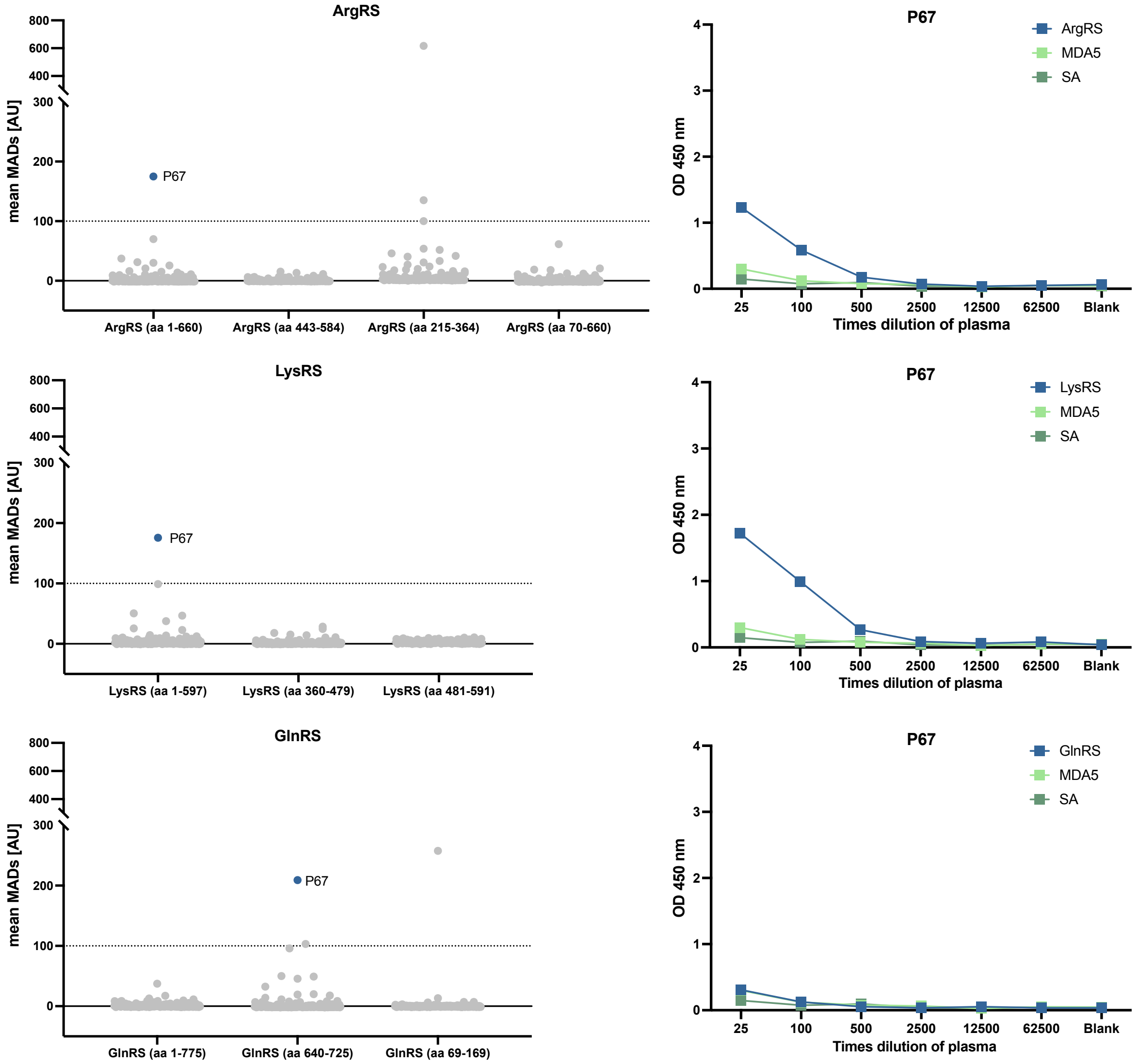


**Supplementary Fig. 7.** Validation of autoantibody reactivities. Results from the multiplex bead array assay (left) and ELISA (right) for P67 targeting three aaRS, all members of the multi-synthetase complex. The amino acid coverage for each antigen used in the multiplex bead array assay is displayed on the x-axis (left). The antigens used in ELISA were ArgRS (aa 1-660), LysRS (aa 1-597), and GlnRS (aa 1-775). aa, amino acid; SA, streptavidin. MDA5, interferon-induced helicase C domain-containing protein 1; MADs, median absolute deviations; OD, optical density.

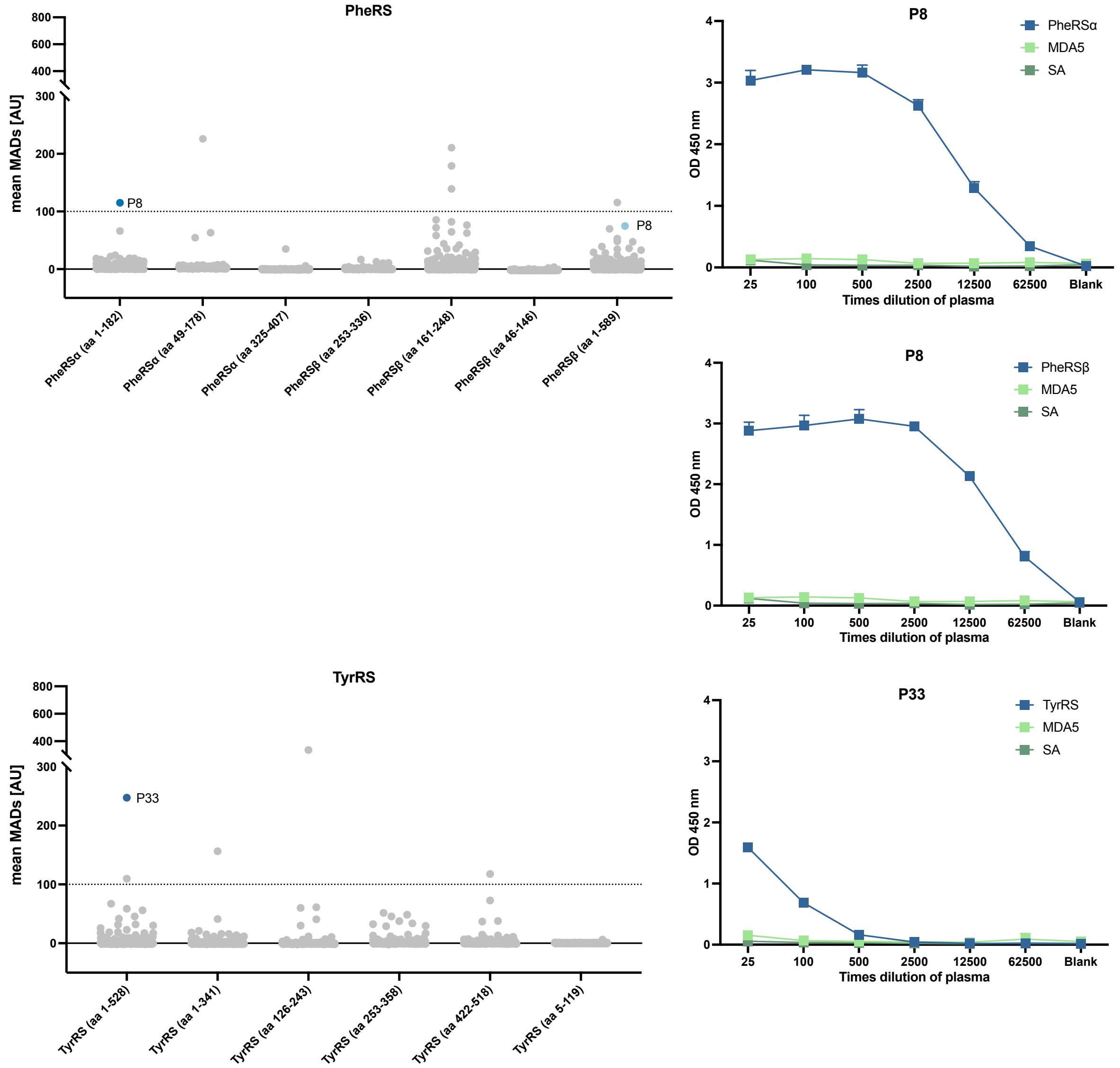


**Supplementary Fig. 8.** Validation of autoantibody reactivities. Results from the multiplex bead array assay (left) and ELISA (right). The amino acid coverage for each antigen used in the multiplex bead array assay is displayed on the x-axis (left). The antigens used in ELISA were PheRSα (aa 1-182), PheRSβ (aa12-589), and TyrRS (aa 1-528). aa, amino acid; SA, streptavidin. MDA5, interferon-induced helicase C domain-containing protein 1; MADs, median absolute deviations; OD, optical density.

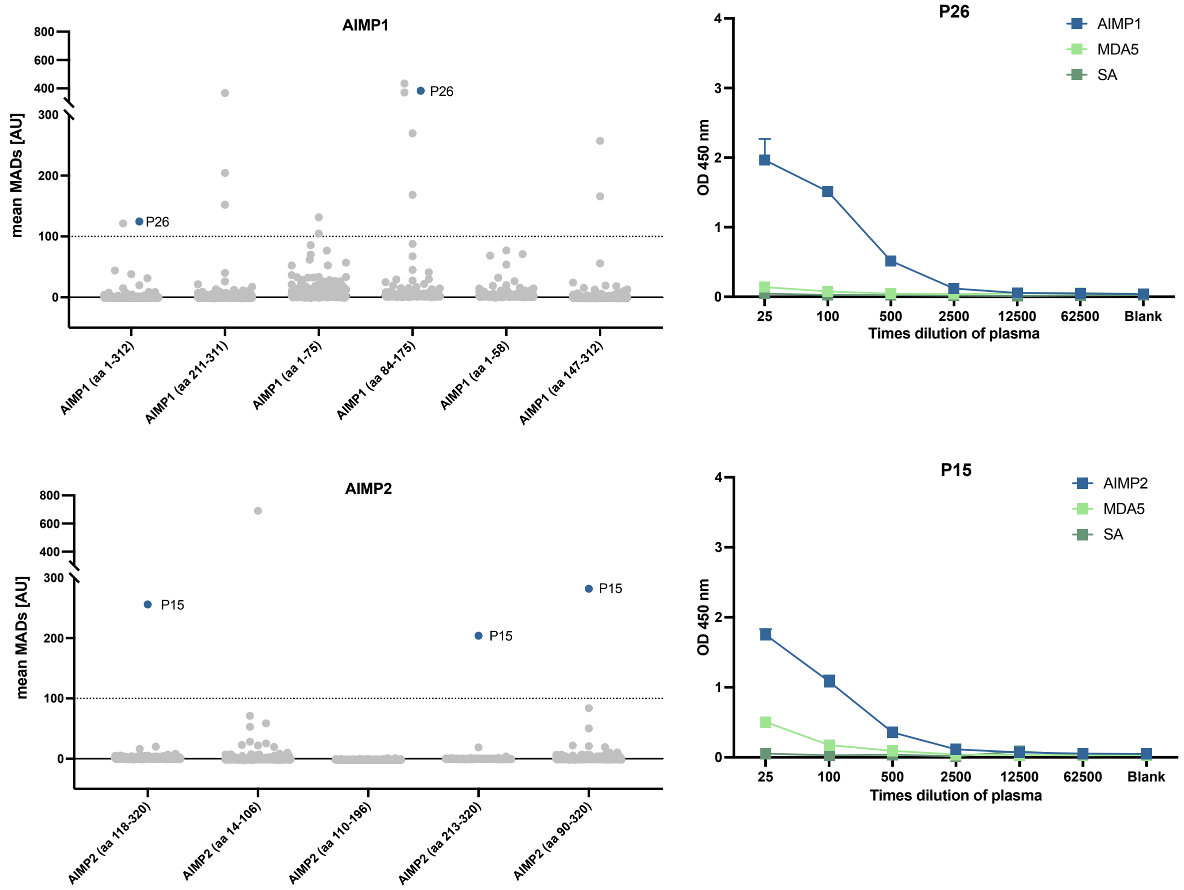


**Supplementary Fig. 9.** Validation of autoantibody reactivities. Results from the multiplex bead array assay (left) and ELISA (right). The amino acid coverage for each antigen used in the multiplex bead array assay is displayed on the x-axis (left). The antigens used in ELISA were AIMP1 (aa 1-312), and AIMP2 (aa 90-320. aa, amino acid; SA, streptavidin. MDA5, interferon-induced helicase C domain-containing protein 1; MADs, median absolute deviations; OD, optical density.

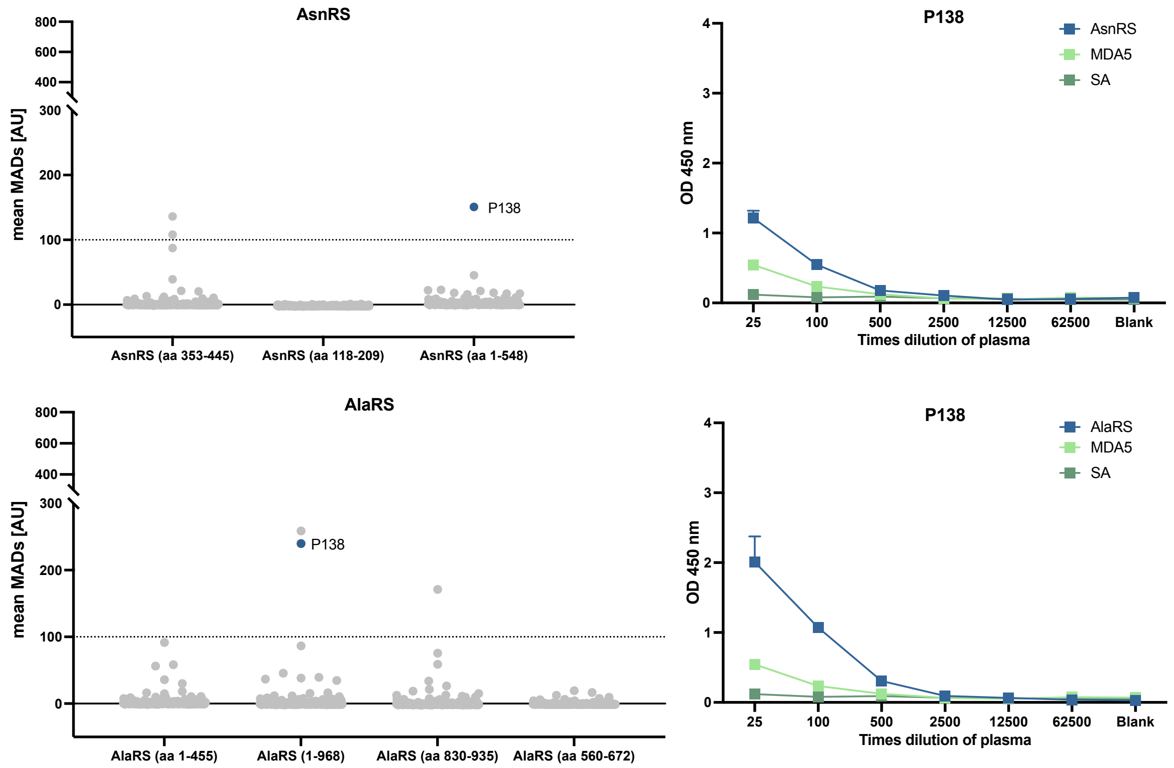


**Supplementary Fig. 10.** Validation of autoantibody reactivities. P138 showing reactivity against both AsnRS (KS) and AlaRS (PL12) in the multiplex bead array assay (left). Both bindings could be validated by ELISA (right). The amino acid coverage for each antigen used in the multiplex bead array assay is displayed on the x-axis (left). The antigen used in ELISA were AsnRS (aa 1-548), and AlaRS (aa 1-968). aa, amino acid; SA, streptavidin. MDA5, interferon-induced helicase C domain-containing protein 1; MADs, median absolute deviations; OD, optical density.

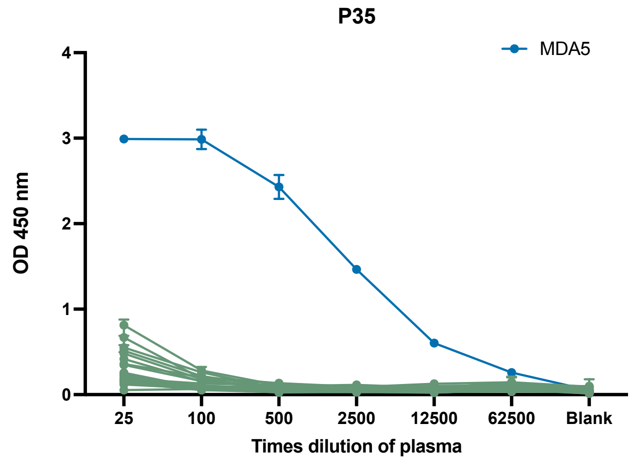


**Supplementary Fig. 11.** P35 (MDA5 positive patient) showing reactivity to MDA5 (aa 110-1025) in blue, but not the other proteins used in ELISA validation (AlaRS, CysRS, PheRSα, PheRSβ, GlyRS, HisRS, AsnRS, SerRS, ThrRS, ValRS, TrpRS, TyrRS, IleRS, ArgRS, GlnRS, LysRS, LeuRS, MetRS, EPRS, AIMP1, and AIMP2) shown in green. MDA5, interferon-induced helicase C domain-containing protein 1; OD, optical density.
